# The epithelial adherens junction component PLEKHA7 regulates ECM remodeling and cell behavior through miRNA-mediated regulation of MMP1 and LOX

**DOI:** 10.1101/2024.05.28.596237

**Authors:** Amanda C. Daulagala, Metin Cetin, Joyce Nair-Menon, Douglas W. Jimenez, Mary Catherine Bridges, Amy D. Bradshaw, Ozgur Sahin, Antonis Kourtidis

## Abstract

Epithelial adherens junctions (AJs) are cell-cell adhesion complexes that are influenced by tissue mechanics, such as those emanating from the extracellular matrix (ECM). Here, we introduce a mechanism whereby epithelial AJs can also regulate the ECM. We show that the AJ component PLEKHA7 regulates levels and activity of the key ECM remodeling components MMP1 and LOX in well-differentiated colon epithelial cells, through the miR-24 and miR-30c miRNAs. PLEKHA7 depletion in epithelial cells results in LOX-dependent ECM remodeling in culture and in the colonic mucosal lamina propria in mice. Furthermore, PLEKHA7-depleted cells exhibit increased migration and invasion rates that are MMP1- and LOX-dependent, and form colonies in 3D cultures that are larger in size and acquire aberrant morphologies in stiffer matrices. These results reveal an AJ-mediated mechanism, through which epithelial cells drive ECM remodeling to modulate their behavior, including acquisition of phenotypes that are hallmarks of conditions such as fibrosis and tumorigenesis.

**Teaser:** Epithelial cells instruct ECM remodeling to modulate their behavior, as a result of adherens junction and miRNA disruption.

## Introduction

Epithelial adherens junctions (AJ) are E-cadherin-based, cell-cell adhesion structures, critical for epithelial barrier integrity and tissue homeostasis (*1, 2*). Intracellularly, E-cadherin binds to p120 catenin (p120) and to β-catenin, and through the latter to α-catenin and the actin cytoskeleton, altogether forming a mechanosensitive complex that regulates cell shape, cell movement, and tissue morphogenesis (*3, 4*). In addition to these well-studied mechanical properties, AJs regulate several signaling pathways in the cell, substantially adding to its role as a hub of cell and tissue homeostasis (*5–7*). Along these lines, we have previously shown that the RNA interference (RNAi) machinery, which is responsible for miRNA biogenesis and function, is recruited to the mature apical epithelial AJs through PLEKHA7, a p120 protein partner (*8–11*). Through this recruitment, PLEKHA7 regulates the levels and activity of a specific set of miRNAs to suppress mRNA expression of proto-oncogenes such as SNAI1, JUN, and MYC (*9–11*). We have shown that loss of PLEKHA7 results in disruption of processing and function of tumor suppressing miRNAs, such as miR-30b, miR- 30c, miR-24, let-7g, miR-200c, resulting in upregulation of their targeting oncogenes, and in increased anchorage independent growth, which is a hallmark of pro-tumorigenic transformation (*9–11*). We have also found that downregulation or loss of PLEKHA7 and disruption of the junctional RNAi machinery is widespread in colon cancer, even in early stages of the disease (*9*).

Importantly, PLEKHA7 depletion neither seems to affect the core E-cadherin-catenin complex, nor to activate β-catenin signaling (*9, 11, 12*). These results showed that PLEKHA7 acts downstream of E-cadherin, corroborating to recent findings underscoring a more nuanced role of the AJs in pro-tumorigenic transformation that does not fall under the classical model of epithelial-to-mesenchymal transition (EMT). In this revised view of the roles of cadherin complexes in tumorigenesis, E-cadherin is still present in tumors and is required for their survival, anchorage-independent growth, and collective cell migration (*9, 11–16*). The findings associating PLEKHA7 with the RNAi machinery, miRNA regulation, and mRNA expression, downstream of E-cadherin have added to this view by incorporating a layer of AJ-mediated regulation of cell behavior through miRNAs.

Here, to comprehensively examine the functions and pathways that PLEKHA7 regulates, we followed an agnostic approach and performed a whole cell RNA-seq analysis in PLEKHA7- depleted cells. Surprisingly, our analysis revealed that the top upregulated pathway in the absence of PLEKHA7 is the one regulating extracellular matrix (ECM) remodeling. Abnormal ECM remodeling, especially increased collagen crosslinking and accumulation, together with fibronectin, is a trait associated with conditions such as cancer and fibrosis (*17–20*). Changes in the ECM can alter adherens junction dynamics, either through actin cytoskeleton-mediated mechanotransduction, or through signaling cascades, and have been extensively documented (*21–25*). Adherens junctions can also influence ECM organization, however, through ways that have been primarily focused on cadherin-integrin-focal adhesion interactions (*26–31*). In this study, we describe a novel mechanism whereby an epithelial adherens junction component influences ECM organization through regulating ECM remodeling enzymes directly, culminating in altered cell behavior.

## Results

### ECM organization is the top PLEKHA7 - regulated cellular pathway

To determine the full breadth of mRNAs and of the downstream cellular pathways regulated by PLEKHA7, we performed whole-cell RNA-seq in the well-differentiated colon epithelial Caco2 cells upon PLEKHA7 depletion and analyzed changes in the mRNA expression profile. Since we have previously found that PLEKHA7 acts as a mRNA suppressor (*9–11*), we focused our analysis on the set of mRNAs that were upregulated upon PLEKHA7 loss. We then ranked the upregulated mRNAs according to the log fold change (Fig. 1A). This ranking identified 1511 mRNAs that their levels were increased upon PLEKHA7 depletion (Fig. 1A and table S1). To narrow down to the potentially most functionally meaningful effects, we selected mRNAs with more than or equal to 1.5 raw fold change. This left 498 mRNAs as the top upregulated ones upon PLEKHA7 depletion (Fig. 1A and table S1). To identify the pathways that are enriched in this set of 498 mRNAs, we performed a pathway analysis using the online Reactome database platform, (v. 84-86) (*32*). In this analysis, we filtered data according to the highest “entities ratio”, the “reaction ratio”, and the lowest p-value. These analyses revealed “extracellular matrix organization” as the top represented pathway in our RNAseq of PLEKHA7-depleted cells (Fig. 1B, and tables S2, S3). We then focused on the specific sub-pathways that are upregulated under the broader ECM organization pathway upon PLEKHA7 depletion. This analysis further revealed “degradation of extracellular matrix” and “collagen formation” as the two most enriched ECM organization sub-pathways in PLEKHA7 - depleted cells, when arranged according to the entities’ ratio (table S4). Together, these results revealed a potential role of PLEKHA7 in ECM organization, which we sought to further investigate.

**Fig. 1.**
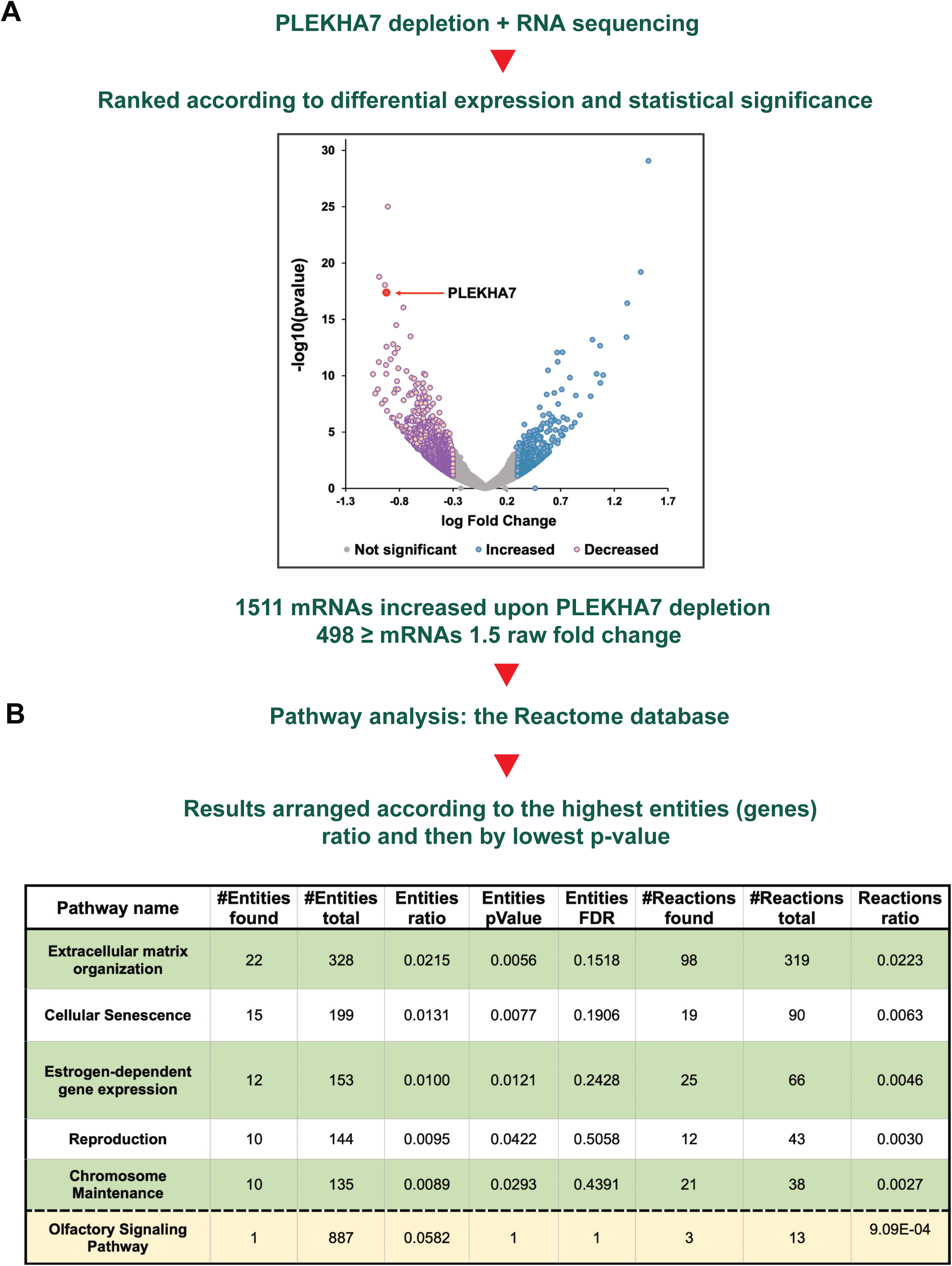
Transcriptomic analysis of PLEKHA7-depleted Caco2 colon epithelial cells reveals ECM as the top upregulated pathway. **(A)** RNA sequencing volcano plot of shRNA - mediated PLEKHA7 knockdown Caco2 cells, compared to non-target shRNA control Caco2 cells from three biological replicates. Results were arranged according to differential log change in expression, to identify mRNAs with the most significant p-values. 1511 mRNAs are increased upon PLEKHA7 depletion, out of which 498 mRNAs exhibit more than or equal to 1.5-fold raw fold change in expression. **(B)** The top upregulated 498 mRNAs were then subjected to pathway analysis against the Reactome database. Results were arranged according to the entities ratio and the significance in p-value, which revealed ECM organization as the most upregulated pathway upon PLEKHA7 depletion. The top five most upregulated pathways are highlighted in green. A pathway highlighted in yellow is shown as a control where only one hit was found, although the pathway includes a large number of total entities in the database, to demonstrate that results are not biased towards pathways with large representation in the database.

### MMP1 and LOX are the top upregulated ECM remodeling hits upon PLEKHA7 depletion

To identify specific ECM-related mRNA hits that are upregulated upon PLEKHA7 depletion, we queried our list of 498 mRNAs against the MatrisomeDB, a database of established ECM markers (*33–35*) (table S5). This analysis revealed 55 ECM - related mRNAs that are upregulated in PLEKHA7 - depleted cells. Combined with the information from the Reactome database analysis, this comparison provided three top upregulated mRNAs that are known regulators of collagen and ECM degradation, namely, meprin 1A, alpha (PABA peptide hydrolase) (MEP1A), matrix metallopeptidase 1 (MMP1), and lysyl oxidase (LOX) (Fig. 2A and table S5). Protein products of all three genes have catalytic activities, where both MEP1A and MMP1 are capable of degrading ECM proteins, while LOX catalyzes collagen crosslinking (*36–41*).

**Fig. 2.**
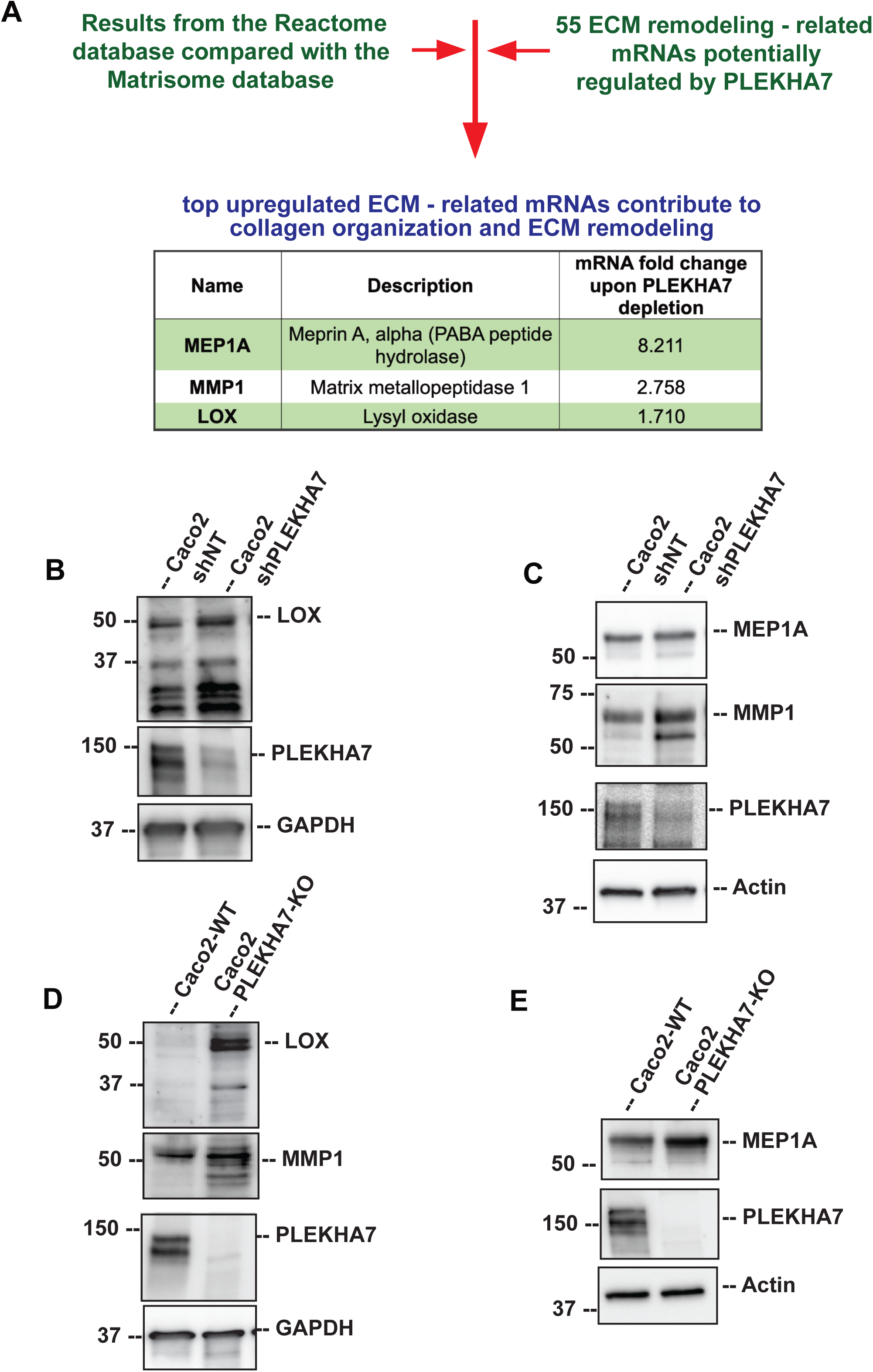
PLEKHA7 depletion upregulates collagen remodeling regulators. **(A)**. The top upregulated 498 mRNAs upon PLEKHA7 depletion were compared against the ECM-specific Matrisome database. This analysis revealed a total of 55 mRNAs that correspond to ECM remodeling. Among these, MEP1A, MMP1 and LOX were the top upregulated mRNAs and matched with the top upregulated ECM organization sub-pathways from the Matrisome analysis, which are “collagen formation” and “matrix degradation” (see also: tables S4 and S5). (**B - E**). Western blot analysis of PLEKHA7, LOX, MMP1, MEP1A, in PLEKHA7 knockdown (shPLEKHA7) and non-target shRNA (shNT), or in wild-type (WT) and PLEKHA7 knockout (PLEKHA7-KO) Caco2 cells. Actin and GAPDH are used as loading controls; molecular weights in kDa are indicated on the left. MMP1 and LOX show consistent upregulation upon PLEKHA7 depletion. Western blots shown are representatives of three biological replicates.

Next, we sought to evaluate whether the increased expression of MEP1A, MMP1 and LOX mRNAs upon PLEKHA7 depletion is also reflected at the protein level. To do this, we performed western blot analysis of cell lysates from Caco2 cells with shRNA-mediated PLEKHA7 knock down (shPLEKHA7), as well as from PLEKHA7 knockout Caco2 cells (PLEKHA7 KO) that we generated using CRISPR/Cas9 (*8*). The results showed that MMP1 and LOX protein levels are indeed consistently increased in both shPLEKHA7 and PLEKHA7 KO cells, compared to the non-target and wild type (WT) Caco2 cells, respectively (Fig. 2B-D). Furthermore, in both cases, it seems that both the pro- (∼47 kDa) and the enzymatically active mature/cleaved LOX (∼37 kDa) and MMP1 (∼54 kDa) protein fragments are increased upon PLEKHA7 depletion (Fig. 2B-E). In contrast, MEP1A protein levels were not changed in shPLEKHA7 cells and only marginally increased in PLEKHA7 KO Caco2 cells and hence did not align with increases in the mRNA levels (Fig. 2A, 2C and 2E). Therefore, we focused our subsequent studies on MMP1 and LOX, both of which showed the most consistent mRNA and protein upregulation upon PLEKHA7 depletion. MMP1 and LOX are master regulators of ECM remodeling.

### MMP1 and LOX are directly and indirectly regulated by PLEKHA7-regulated miRNAs

We have previously shown that PLEKHA7 suppresses mRNA levels through a set of miRNAs (*9–11*). Therefore, we then examined whether PLEKHA7 suppresses MMP1 and LOX mRNA levels through its regulated miRNAs. First, we used the bioinformatic miRNA target prediction tools DIANA (version 8.0) (*42*), mirTarBase (release 8.0) (*43*), and TargetScan (release 7.2 and 8.0) (*44, 45*), to identify which PLEKHA7-regulated miRNAs from our published datasets (*10, 11*) could be targeting MMP1 and LOX. This analysis identified miR-203a, let-7g, miR-30b, miR-30c, and miR- 24 as potentially targeting the MMP1 and LOX mRNAs (Fig. 3A). We have shown that PLEKHA7 depletion results in downregulation of these miRNAs (*9–11*). Thus, we wanted to examine if the increased protein levels of MMP1 and LOX can be rescued using miRNA mimics of these miRNAs that are predicted to target the mRNAs of these markers. To do this, we transfected miR-203a, let- 7g, miR-24, miR-30b and miR-30c mimics into PLEKHA7 KO Caco2 cells. Western blot analysis revealed that miR-24 and miR-30c mimics were able to rescue the increased protein levels of MMP1 and LOX upon PLEKHA7 knockout (Fig. 3B and 3C). miR-24 and miR-30c are indeed downregulated in PLEKHA7 KO cells (Fig. 3D).

**Fig. 3.**
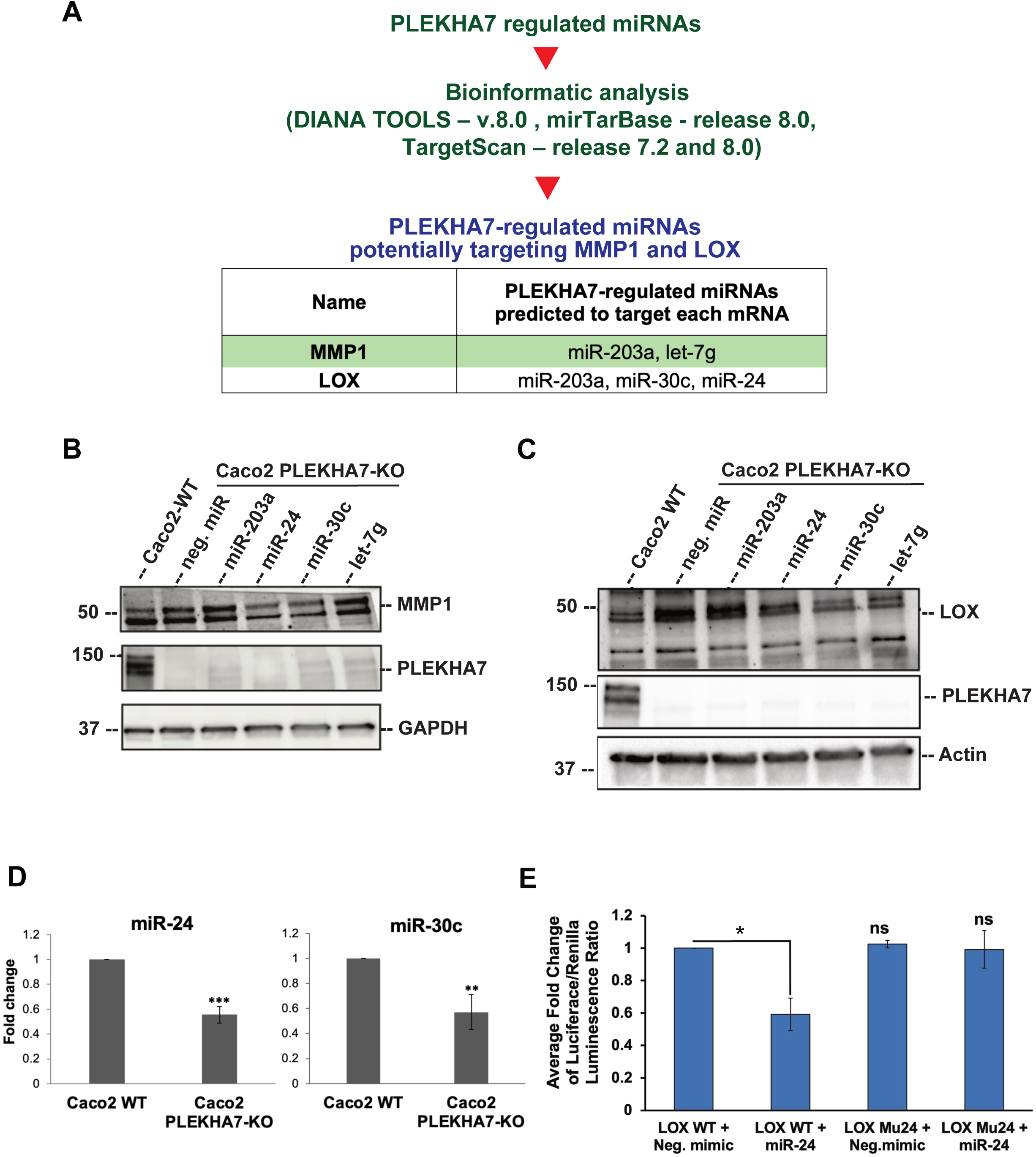
PLEKHA7 regulates MMP1 and LOX through miRNAs. **(A)** Bioinformatic analysis of a previously identified set of PLEKHA7-regulated miRNAs using DIANA TOOLS (v.8.0), mirTarBase (release 8.0), and TargetScan (release 7.2 and 8.0), to identify whether they are predicted to target MMP1 and LOX mRNAs. This analysis revealed miR-203a and let-7g as potentially targeting the MMP1 mRNA, and miR-24, miR-30c, and miR-203a as potentially targeting the LOX mRNA. (**B, C**) Western blot analysis of Caco2-WT and PLEKHA7-KO cells transfected with either negative control miRNA mimic or miRNA mimics of the above miRNAs, to examine MMP1 and LOX rescue in expression. Western blot of PLEKHA7 indicates knockout and GAPDH, Actin are used as loading controls; molecular weights in kDa are indicated on the left. (**D**) qRT-PCR analysis of Caco2-WT or PLEKHA7-KO cells for miR-24 and miR-30c to confirm their downregulation upon PLEKHA7 depletion. ***p=0.0003; **p=0.005, student’s t-test, n=3 biological replicates. (**E**) Luciferase reporter assay using LOX 3’UTR luciferase constructs, wild type (LOX WT) and mutated for the miR-24 target sequence (LOX Mu24), transfected in PLEKHA7 KO Caco2 cells and co-transfected with negative control miRNA mimic, or miR-24 mimic. Results shown are the average luciferase/renilla luminescence ratio from n=3 biological replicates; p=0.05, one-way ANOVA with Bonferroni correction.

miRNA target sequences are typically at the 3’ untranslated regions (3’UTRs) of mRNAs. Our miRNA prediction analysis identified target sites for both miR-24 and miR-30c on the LOX mRNA 3’UTR. To verify that the miR-24 and miR-30c effects on LOX are through direct targeting of its mRNA 3’UTR, we generated luciferase reporter constructs bearing the wild-type 3’UTR target region of LOX mRNA, as well as the 3’UTR target region mutated, to disrupt binding of miR-24 and miR-30c through their seed sequence. We then transfected Caco2 cells with these LOX 3’UTR luciferase reporter constructs, together with miR-24 and miR-30c mimics, to measure mRNA degradation through miRNA targeting. Results indicate that expression of the LOX mRNA is indeed inhibited by miR-24, whereas this inhibition is rescued when the miR-24-targeting sequence on the LOX mRNA is mutated (Fig. 3E). However, there was no noticeable inhibition even of the wild-type LOX mRNA 3’UTR by miR-30c (fig. S1). Thus, our results show that the LOX mRNA is directly targeted by miR-24, but not by miR-30c.

Although miR-30c does not seem to target the LOX mRNA 3’UTR directly, our western blot results showed that the miR-30c mimic does rescue LOX protein levels in PLEKHA7-KO cells (Fig. 3C). In addition, both miR-24 and miR-30c rescue MMP1 protein levels upon PLEKHA7 depletion (Fig. 3C and 3D), although our bioinformatic analysis did not identify direct seed sequence binding of either miR-24 or miR-30c to the 3’UTR of the MMP1 mRNA. These observations indicate that PLEKHA7 regulates LOX through miR-30c and MMP1 through miR-24 or miR-30c not directly, but through intermediate targets. One such intermediate target could be the JUN (cellular homolog of Jun avian sarcoma virus 17 proto-oncogene), which is responsible for MMP1 and LOX mRNA transcription (*46–49*) and that we have previously shown to be suppressed by PLEKHA7 through miR-24 (*10, 11*). Additional studies have also shown that miR-30 miRNAs suppress JUN (*50*). Based on the above, we examined potential regulation of MMP1 by miR-24 and miR-30c via JUN. First, we confirmed increased protein levels of JUN in PLEKHA7 KO cells, which are rescued by miR-24 and miR-30c mimics, demonstrating JUN regulation by miR-24 and miR-30c (Fig. 4A). Then, to examine whether JUN is indeed the intermediate of PLEKHA7 - regulation of MMP1 and LOX, we assessed MMP1 and LOX protein levels in PLEKHA7 KO cells by simultaneously performing shRNA-mediated knockdown of JUN (Fig. 4B and 4C). Although cells could not tolerate complete downregulation of JUN, since it is an essential transcription factor and we could only achieve moderate levels of JUN knockdown, that was sufficient to rescue MMP1 and LOX levels in PLEKHA7 KO cells to their baseline levels (Fig. 4B and 4C). Together, these results outline a pathway where PLEKHA7 regulates LOX mRNA directly, via miR-24, and MMP1 and LOX indirectly, by targeting JUN via miR-24 and miR-30c (Fig. 4D).

**Fig. 4.**
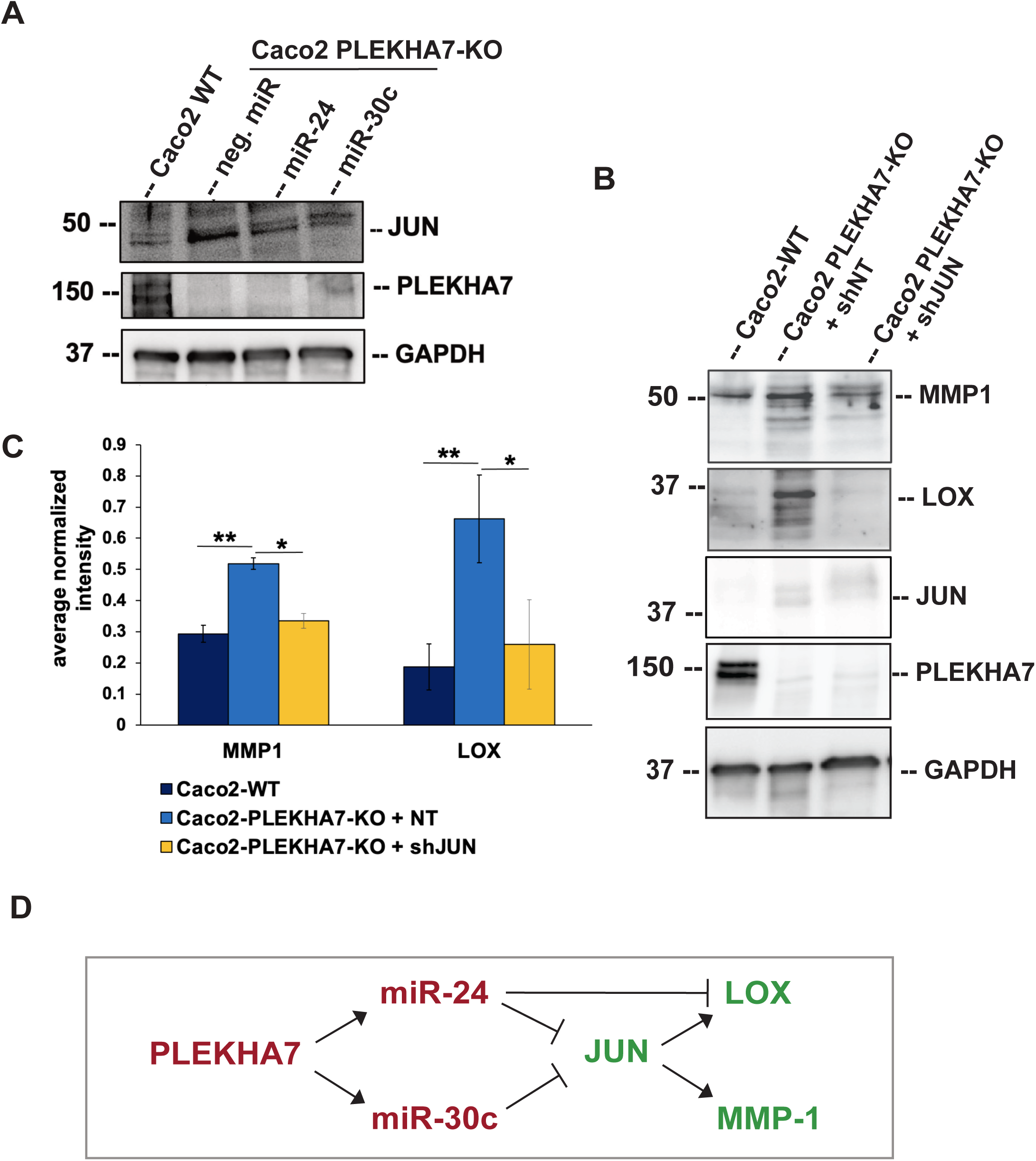
PLEKHA7 regulates MMP1 and LOX also through miRNA-mediated regulation of JUN. **(A)**. Western blot of Caco2-WT and PLEKHA7-KO cells transfected with miR-24 and miR- 30c mimics for JUN and PLEKHA7; GAPDH is the loading control; molecular weights in kDa are indicated on the left. **(B)** Western blot of Caco2-WT and PLEKHA7-KO cells transduced with non- target (NT) or JUN shRNA, for MMP1, LOX, PLEKHA7, JUN; GAPDH is the loading control; molecular weights in kDa are indicated on the left. **(C)** Quantification of the western blot from (B) for MMP1 and LOX protein levels, normalized to GAPDH and averaged from n=3 biological replicates; *p<0.05, **p<0.01, one-way ANOVA with Bonferroni correction. **(D)** Summary of results from Figs 1-4, showing that PLEKHA7 regulates LOX directly through miR-24, and both MMP-1 and LOX through miR-24 and miR-30c - mediated regulation of JUN.

### PLEKHA7 knockout results in increased MMP1 and LOX activities and ECM remodeling

MMP1 is a collagenase that acts on collagen I as its substrate, while LOX is an oxidase that catalyzes the formation of covalent crosslinks in collagen fibrils and fibers (*38, 39, 51*). Therefore, we next sought to examine whether the increase in MMP1 and LOX protein levels is also accompanied by increased catalytic activity of these enzymes. To assess this, we employed collagenase and LOX activity assays, using commercially available, fluorescence-based kits. Indeed, these activity assays showed significantly increased MMP1 and LOX activities in PLEKHA7 KO Caco2 cells (Fig. 5A and 5B).

**Fig. 5.**
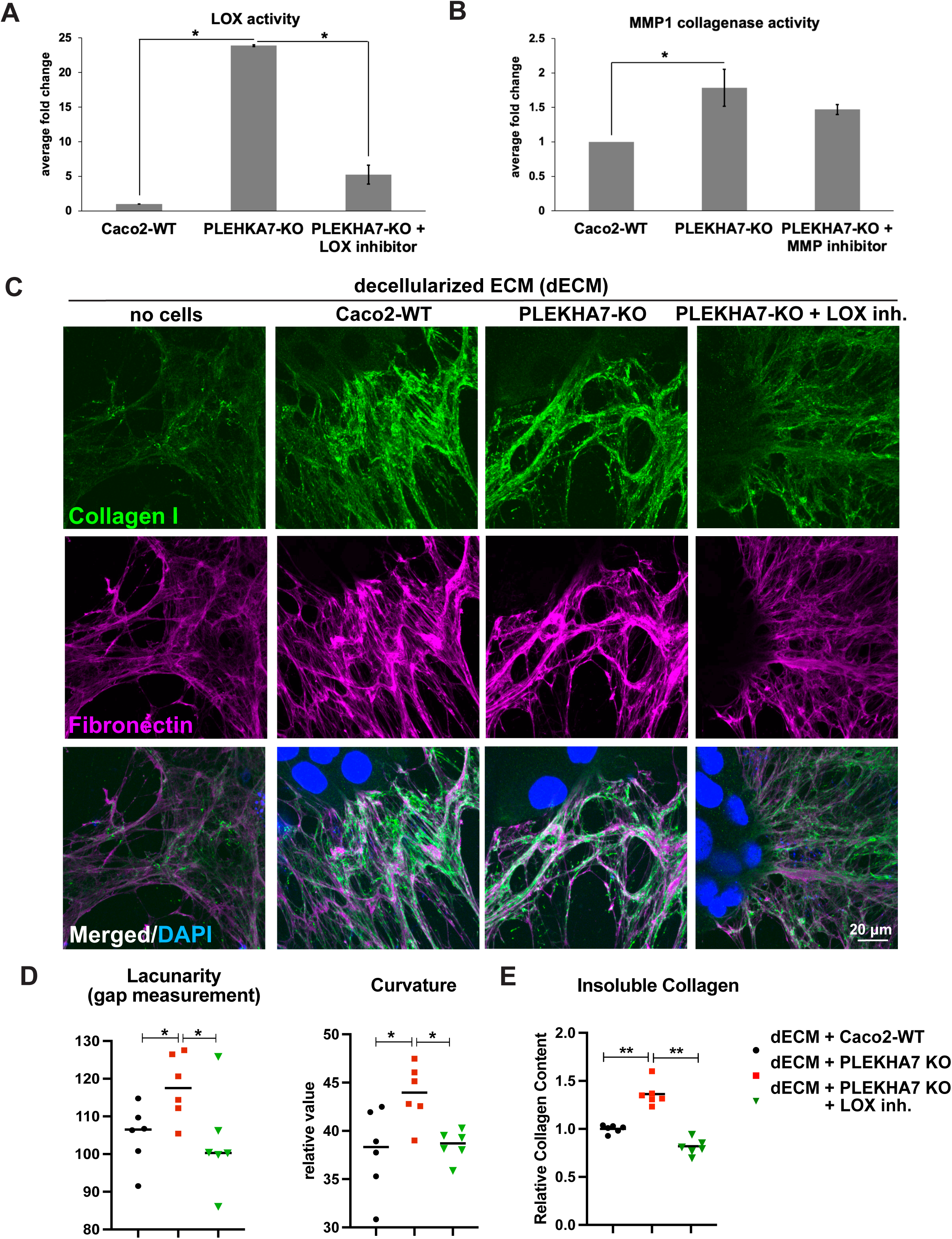
PLEKHA7 depletion promotes LOX and MMP1 activities and ECM remodeling. **(A)** Measurements of LOX and **(B)** MMP1 enzymatic activities in Caco2-WT and PLEKHA7-KO cells, and after treatment of PLEKHA7-KO cells with LOX (BAPN; 150 µM final) and MMP (Doxycycline Hyclate; 0.2 µg/mL final) inhibitors, respectively. **(C)** Immunostaining of collagen I and fibronectin of HFF-1-derived decellularized ECM and upon plating it with Caco2-WT, PLEKHA7-KO cells, and PLEKHA7-KO cells treated with the LOX inhibitor BAPN, as above. **(D)** Quantification of images from C, using the TWOMBLI macro on Fiji (*55*), from n=6 fields representative of two biological replicates; *p<0.05, one-way ANOVA. **(E)** Total collagen assay for the insoluble crosslinked-collagen fraction of Caco2-WT, PLEKHA7-KO cells, and PLEKHA7-KO cells treated with the LOX inhibitor BAPN, as above, from n=2 biological replicates; **p<0.01, one-way ANOVA.

The results showing increased MMP1 and LOX activity levels upon PLEKHA7 depletion in the epithelial Caco2 cells suggest effects on ECM remodeling driven by these cells. Since the primary source of ECM in tissues is typically fibroblasts, we cultured human foreskin fibroblast cells (HFF-1) to produce ECM. Then, we de-cellularized the ECM from the HFF-1 fibroblasts and plated WT and PLEKHA7 KO Caco2 cells on their produced ECM. Examination of collagen I as well as of fibronectin that is also critical for collagen I matrix organization (*52, 53*) by immunofluorescence, showed that compared to the de-cellularized ECM, Caco2-WT cells seem to effectively organize the ECM in a homogeneous, isotropic matrix (Fig. 5C). However, PLEKHA7 KO cells seem to further remodel these matrices in better organized and thicker bundles, resulting in more anisotropic ECM compared to the WT cells (Fig. 5C). This was evident by higher lacunarity, or else the presence of larger gaps between fibril bundles, and higher curvature (Fig. 5D), both typical of anisotropic matrices (*54*). Strikingly, this effect of the PLEKHA7 KO cells on the ECM was reversed to resemble the one organized by the Caco2-WT cells upon treatment with BAPN (β-aminopropionitrile), an inhibitor of LOX, showing that the observed PLEKHA7 KO-mediated ECM remodeling is LOX-dependent. This was further supported biochemically using a total collagen assay to quantify crosslinked collagen via pepsin digested insoluble collagen I fraction. This assay showed that insoluble collagen I was significantly elevated in the matrix plated with PLEKHA7 KO cells compared to the matrix with Caco2-WT cells, a result that was again reversed upon treatment with the LOX inhibitor (Fig. 5E), further demonstrating that the effects of PLEKHA7 depletion on the ECM are LOX-dependent. Altogether, these results demonstrate that PLEKHA7 depletion in colon epithelial Caco2 cells results in increased MMP1 and LOX activities, which in turn promote ECM remodeling driven by the PLEKHA7-depleted epithelial cells.

### PLEKHA7 depletion promotes MMP1- and LOX-dependent cell migration and invasion

Increased MMP1 and LOX activities and aberrant ECM remodeling are characteristics of wound healing and invasion (*55, 56*). ECM remodeling may promote cell mobility, whereas LOX and MMP1 upregulation in tumors has been extensively implicated in cancer cell invasion and metastasis (*57, 58*). Indeed, both MMP1 and LOX are crucial for collagen fibril formation, whereas increased collagen fiber re-organization is a common occurrence in the tumor microenvironment (*59*). Thus, we sought to investigate if the increased MMP1 and LOX activities observed upon PLEKHA7 depletion also promote epithelial cell migration and invasion. To do this, we plated Caco2 cells on Matrigel-coated coverslips to perform a wound healing assay and measured the wound closure area over time. We limited our study duration to <32 hours, to avoid any effects due to cell proliferation, since Caco2 cells have a doubling time of 32 hours (*60*). This experiment showed that PLEKHA7 depletion results in increased wound closing rates compared to control Caco2 cells (Fig. 6A and 6B). We then examined cell invasion rates in real time, by using the xCELLigence Real-Time Cell Analysis (RTCA) system, where we plated cells on collagen I - coated - RTCA Cell Invasion (CIM) plates. The results showed that PLEKHA7 KO cells exhibit significantly increased invasion rates, compared to the Caco2-WT cells, which are in essence non-invasive, as expected for these well-differentiated epithelial cells (Fig. 6C and 6D). To examine whether the invasion abilities of PLEKHA7 KO cells are due to the increased MMP1 and LOX activities, we used the MMP inhibitor Doxycycline hyclate and the LOX inhibitor BAPN and measured the respective invasion rates of PLEKHA7-depleted cells using the same assay. Indeed, the increased invasion rates of PLEKHA7 KO cells were rescued upon either MMP1 or LOX inhibition, indicating that the observed ability of Caco2 cells to invade upon loss of PLEKHA7 is due to the increased MMP1 and LOX activities (Fig. 6C and 6D). According to the manufacturer’s specifications (Santa Cruz), Doxycycline hyclate may inhibit MMP1, MMP8, and MMP9; however, in our RNA-seq data only MMP1 is significantly upregulated upon PLEKHA7 depletion, whereas MMP8 and MMP9 are not expressed in these cells (table S6). Similarly, although BAPN could potentially inhibit other LOX family members, these either did not increase by PLEKHA7 depletion, or are not expressed at significant levels in these cells (table S6). Therefore, rescuing of the increased invasion rates that we observed upon PLEKHA7 depletion are specifically mediated by the upregulation of MMP1 and LOX activities in the PLEKHA7-depleted cells.

**Fig. 6.**
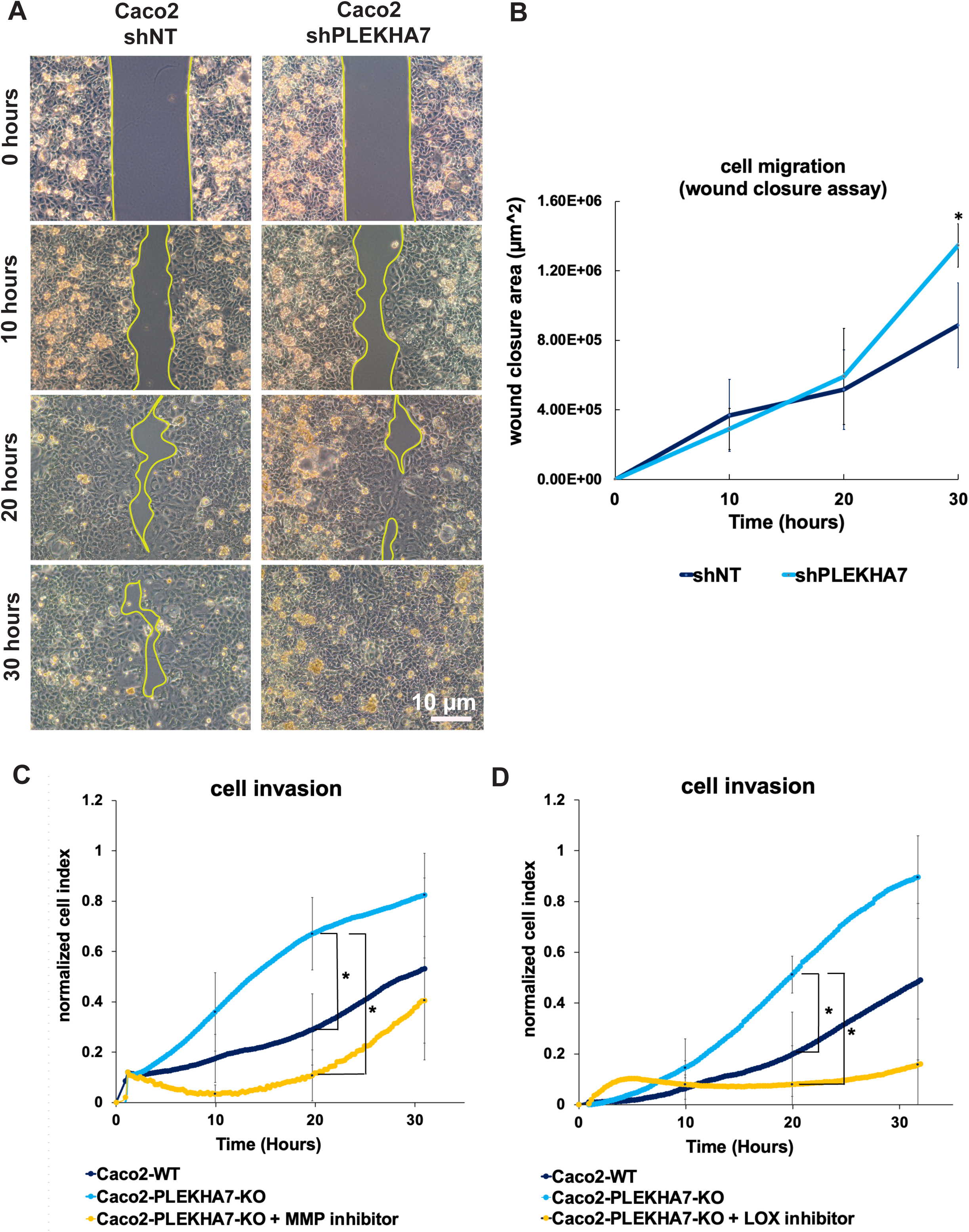
PLEKHA7 depletion promotes cell migration and invasion. **(A)** Caco2 with non-target shRNA (shNT) and with shRNA mediated PLEKHA7 knockdown (shPLEKHA7), were plated on Matrigel - coated coverslips using Ibidi 2-well cell inserts till full confluency. Once confluent, inserts were removed and cell growth into the “wound” area (indicated within yellow lines) was imaged every 10 hours and **(B)** quantified **(\***p=0.04, from n=2 biological replicates, Student’s t-test). **(C, D)** Caco2-WT, PLEKHA7-KO, and PLEKHA7-KO cells treated with the MMP inhibitor (Doxycycline hyclate: 0.2 µg/mL) or the LOX inhibitor (BAPN: 150 µm), were plated on CIM transwell plates and cell invasion was measured in real time using the xCELLigence real time cell analysis system. All wound closure and invasion experiments were stopped at <32 hours to avoid effects of proliferation. *p<0.05, from n=3 biological replicates, one-way ANOVA with Bonferroni correction.

### PLEKHA7 knockout cells grow abnormally in 3D cultures and in stiffer matrices

Our results indicate that PLEKHA7 depletion promotes ECM remodeling, as well as in increased cell migration and invasion rates, though increasing MMP1 and LOX activities. Since these assays were performed in 2D cultures, we also investigated any phenotypic changes in PLEKHA7 depleted cells when these cells are grown in 3D cultures. To do this, we plated Caco2 WT and PLEKHA7 KO cells in low stiffness (1.1 kPa) and in denser, higher stiffness (3 kPa) 3D matrices using the RASTRUM platform (*61, 62*). Our results show that PLEKHA7 KO cells grow larger and lose circularity compared to the Caco2 WT cells, when embedded in normal matrices (Fig. 7A, 7B and fig. S2). Interestingly, in the tumor-mimicking matrices, even Caco2 WT cells grew larger and lose circularity, compared to Caco2 WT cells grown in normal matrices (Fig. 7A,7B and fig. S2). In fact, both Caco2 WT in the stiffer (3 kPa) matrix and PLEKHA7 KO cells in the softer matrix (1.1 kPa) exhibit similar attributes in terms of area and circularity of the spheroids (Fig. 7B). Although PLEKHA7 KO cells overall retained their circular structure in the normal matrix, they significantly lost circularity and grew in larger disorganized formations in the tumor mimicking matrix (Fig. 7A, 7B and fig. S2), which is reminiscent of pro-tumorigenic transformation (*63, 64*). We obtained similar results when we plated Caco2 cells with and without PLEKHA7 in commercially available hydrogels (VitroGel), which also mimic normal (softer) and tumor (stiffer) conditions (fig. S3).

**Fig. 7.**
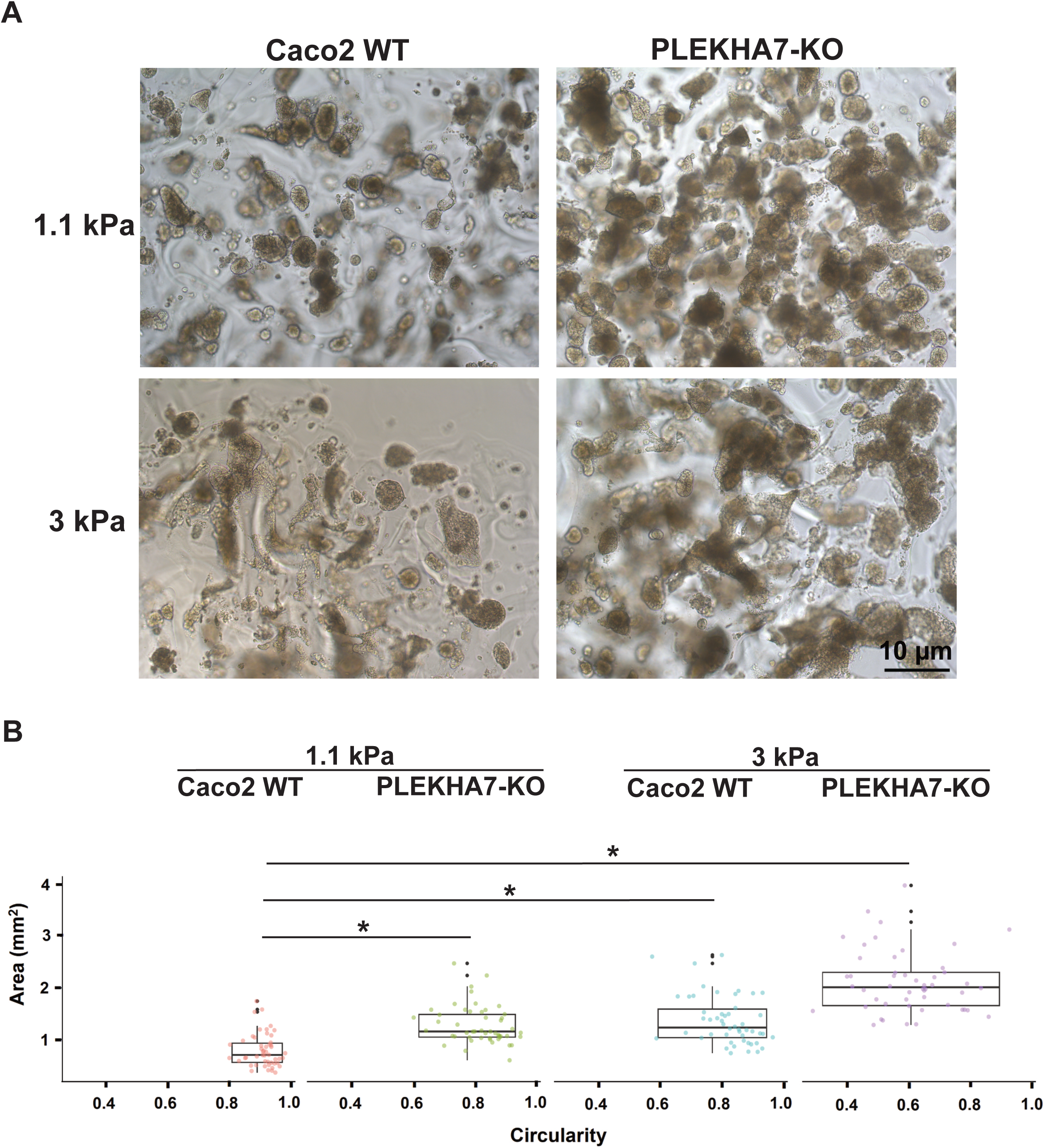
PLEKHA7 knockout promotes aberrant growth of Caco2 cells in 3D cultures when combined with a stiff ECM substrate. **(A, B)** Caco2-WT and PLEKHA7-KO cells were grown in 3D cultures in 1.1 kPa and in stiffer, 3 kPa matrices, using a RASTRUM apparatus for 10 days and then imaged and quantified for size and for circularity, to assess shape. Asterisk indicates the represented condition is significantly different compared to the Caco2-WT cells in the softer 1.1 kPa matrix, which was used as the reference condition. *p<0.05, from n=50 colonies representative of two biological replicates, two-way ANOVA with Bonferroni correction.

Overall, these results indicate that PLEKHA7 depletion promotes transformation of colon epithelial cells in 3D and in coordination with the ECM microenvironment.

### *Plekha7*^-/-^ mice exhibit increased collagen deposition in the colonic lamina propria

Our in vitro studies showed that PLEKHA7 depletion in Caco2 colon epithelial cells results in collagen matrix remodeling through upregulation of MMP1 and LOX, in vitro. We then wanted to examine whether we would observe any effects on collagen remodeling by PLEKHA7 loss, in vivo. To do this, we generated a *Plekha7* knockout mouse model on C57BL/6J background using CRISPR/Cas9 technology. Since multiple Plekha7 isoforms are potentially expressed in mice, we deleted exons 2-10 of *Plekha7* spanning 30 kb and through using two gRNAs, to ensure complete gene knockout (fig. S4A). Examination of the UCSC genome browser (mm39) does not show any other genes expressed from that region that their loss would introduce any non-specific effects (fig. S4B). Plekha7 is specifically expressed in the epithelium, at the apical areas of cell-cell contact and in the colonic epithelial crypts, as we and others have previously shown (*9, 11, 65*) and which we also confirmed here by colon tissue immunofluorescence (Fig. 8A). These mice are overall viable, and no developmental defects were observed.

**Fig. 8.**
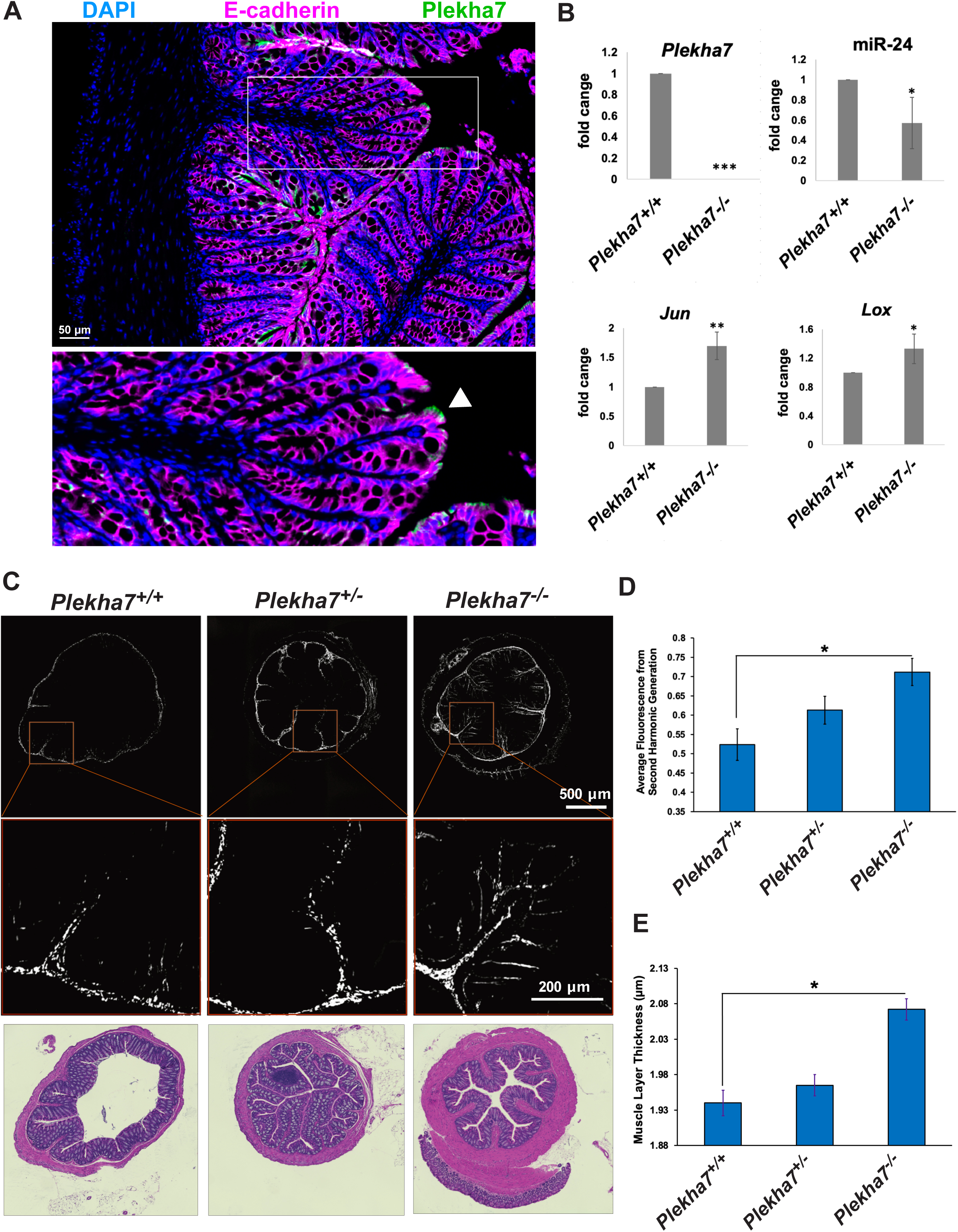
PLEKHA7 knockout results in increase collage deposition in the colonic mucosa, in vivo. **(A)** Immunofluorescence of wild type mouse colon tissue for Plekha7 and E-cadherin; DAPI is the nuclear stain. An inset is shown to the right; arrowhead indicates apical epithelial Plekha7 localization. **(B)** qRT-PCR analysis of *Plekha7^+/+^* and *Plekha7^-/-^*mouse whole tissue colon RNAs for *Plekha7* mRNA, miR-24 (mmu-miR-24), *Jun* and *Lox* mRNAs. ***p<0.005; **p<0.01, *p<0.05, student’s t-test, n=3 mice. **(C)** Second harmonic generation images (top row) and representative enlarged insets (middle row) of colon sections of six-month-old *Plekha7^+/+^*, *Plekha7^+/-^,* and *Plekha7^-/-^* mice were obtained to analyze collagen deposition. Bottom row shows the corresponding H&E images of the same sections. **(D, E)**. Second harmonic generation (collagen) and muscle layer thickness quantifications from (B) from n=25 mice (distribution: *Plekha7^+/+^*= 9, *Plekha7^+/-^* = 8, *Plekha7^-/-^* = 8; Female = 13, Male = 12). *p<0.05, one-way ANOVA with Bonferroni correction.

We confirmed complete *Plekha7* knockout by qRT-PCR in colon tissues from *Plekha7*^-/-^ mice (Fig. 8B). Furthermore, and corroborating with our human Caco2 studies, the *Plekha7*^-/-^ colon tissues show significant downregulation of miR-24 and significant increase in expression levels of *Jun* and *Lox* (Fig. 8B). We were not able to detect *Mmp1* in these tissues, when examining either of the two mouse *Mmp1a* or *Mmp1b* variants, since *Mmp1* is overall not expressed in the normal mouse colon (*66*). However, since *Lox* was upregulated and since our in vitro data showed LOX-dependent collagen I remodeling upon PLEKHA7 loss in human colon epithelial cells, we then examined collagen organization in the colon tissues of *Plekha7*^-/-^ mice. To do this, we imaged colon tissues of 25 six-month-old mice using second harmonic generation microscopy to visualize and quantify fibrillar collagen (Fig. 8C). This approach showed that fibrillar collagens in the colonic lamina propria of *Plekha7*^-/-^ mice are significantly increased compared to *Plekha7*^+/+^ (Fig. 8C and 8D), in line with the increased expression of LOX. Interestingly, although collagen is almost explicitly found in the submucosa of the *Plekha7*^+/+^ mice, collagen deposition was seen to be significantly expanded in the mucosa of the *Plekha7*^+/-^ mice, and even in the mucosal lamina propria of the of the *Plekha7*^-/-^ mice (Fig. 8C). Increased collagen deposition in the colon is a key characteristic of fibrosis and can promote tumorigenesis (*67–69*). Strikingly, the overall muscular layer of *Plekha7*^-/-^ mice was significantly thickened, which is another hallmark of fibrosis in the colon (Fig. 8C and 8E)(*69–71*). Altogether, our results using *Plekha7* knockout mice corroborate our cell culture experiments and further support our in vitro findings revealing that the AJ component PLEKHA7 regulates ECM remodeling in epithelial cells and tissues.

## Discussion

The colon epithelial barrier is a focal point of investigation, due to its central role in diseases of increasing prevalence, especially among the younger population, such as inflammatory bowel disease, fibrosis, and colon cancer (*72–74*). AJs play a crucial role in maintaining epithelial barrier integrity and their disruption has been consistently associated with these diseases (*2, 75*). Indeed, the epithelial compartment is where most colon tumors originate from (*76, 77*). Epithelial cells attach to the ECM through focal adhesions, which in turn communicate with the AJs via the integrin-cytoskeleton network (*78, 79*). Thus, alterations in the ECM can be sensed by AJs and vice versa. Along these lines, several studies have revealed signaling or biophysical ways through which the AJs and ECM communicate with each other, to regulate cellular functions, such as cell migration (*13, 80–83*).

Our findings add a new layer of regulation in the AJs-ECM crosstalk, by revealing a mechanism that links the epithelial AJ component PLEKHA7 that is critical for AJ integrity and barrier function (*8, 11*), with regulation of ECM remodeling. Although we have previously shown that PLEKHA7 recruits and regulates the RNAi machinery and miRNAs to the AJs to suppress a number of oncogenes, we had only identified a specific set of PLEKHA7’s targets, primarily guided through the predicted targets of the PLEKHA7-associated miRNAs (*9–11*). In this work, through interrogating the whole set of potential downstream targets of PLEKHA7 by employing RNAseq and extensive bioinformatic analyses, we identified ECM regulation as being the top cellular function regulated by PLEKHA7. Furthermore, we specifically found that PLEKHA7 depletion increases the levels of a set of ECM remodeling markers involved in collagen fibril formation and degradation, such as MMP1 and LOX. The increased mRNA and protein levels of MMP1 and LOX upon PLEKHA7 depletion in Caco2 epithelial cells also result in elevated activity levels of MMP1 and LOX (Fig. 5A and 5B) and affects fibroblast-produced ECM remodeling (Fig. 5C-E). This is striking, since ECM remodeling is generally driven by fibroblasts’ activity, but not of that of epithelial cells (*84, 85*). Moreover, although we focused our experimentation on MMP1 and LOX that are two master ECM regulators, our RNA-seq and bioinformatic analysis revealed a broader set of additional ECM components and regulators, such as Fibrinogen, Cystatin SN and SPOCK3 (Fig. 1 and table S5). These findings reveal a whole program of ECM regulation by PLEKHA7, which deems to be further investigated with regards to its extent and the specific roles of each one of these components in the observed ECM remodeling phenotypes.

ECM remodeling mediated by increased LOX and MMP1 activities is a main occurrence in events such as cell migration during wound healing and cell invasion (*55, 56*). Since PLEKHA7 depletion results in ECM remodeling through LOX and MMP1 upregulation, we hypothesized that this will also promote wound closure and cell invasion. Indeed, although Caco2 cells are well-differentiated and overall non-invasive (Fig. 6A-D), PLEKHA7 depletion accelerates wound closing and induces collagen matrix invasion in these cells (Fig. 6A-D). Although faster wound closure rates could be beneficial in normal physiological conditions, a continuous wound closure and epithelial cell-mediated invasion can lead to abnormal cell behavior, such as a constantly dividing cells mass, which is a characteristic of tumors, as well as scar tissue formation that is observed in fibrosis. Furthermore, PLEKHA7-depleted Caco2 cells grow in larger spheroids when cultured in 3D matrices and lose their spherical shape in stiffer, tumor-mimicking matrices, compared to the wild type cells, which overall retained a spherical shape in either condition (Fig. 7A and B). This observation is reminiscent of the “second hit hypothesis” and the “seed and soil theory”; according to these ideas, PLEKHA7 loss is the first hit creating bad “seeds” that could be precancerous, whereas the eventual ECM remodeling that occurs to favor the growth of PLEKHA7 depleted cells is the “soil” and the second hit. Supporting this notion and our in vitro studies, *Plekha7^-/-^* mice exhibit increased collagen deposition in the mucosal lamina propria in the colon compared to the *Plekha7^+/+^*mice, which is a characteristic of both fibrotic and tumor conditions (Fig. 8C and D). Although we did not observe tumor formation or cell invasion in our *Plekha7^-/-^* mice, this is likely due to the fact that additional “hits” are likely required to observe such a phenotype. Indeed, the vast majority of colon tumors arise from mutations on the *APC* tumor suppressor gene, whereas additional hits, such as *P53* and *KRAS* mutations, are required for colon cancer development (*86, 87*). Caco2 cells do harbor *APC* and *P53* mutations (*88, 89*), which is potentially why they exhibit pro-tumorigenic phenotypes upon PLEKHA7 depletion (Fig. 6A-D and 7A-B). It is likely that events such AJ-mediated dysregulation of the ECM provide an initial “hit”, which predisposes to colon tumorigenesis in the event of additional mutations. Fibrosis and increased collagen I accumulation are hallmarks of inflammatory bowel disease (IBD), which strongly increases colon cancer risk (*90*). In this context, it would be interesting to examine in our upcoming studies whether PLEKHA7-mediated disruption of the ECM can predispose colon tissues to tumorigenesis, both in mice and in human patients.

Our results show that PLEKHA7 suppresses MMP1 and LOX through its associated miR- 24 and miR-30c miRNAs (Fig. 3C and 3E). We have previously shown that PLEKHA7 depletion results in loss of junctional localization of AGO2, which is the key enzymatic component of the RNAi machinery, as well as in decreased miRNA loading to AGO2. We have also shown that PLEKHA7 is not the only AJ component that mediates AGO2 recruitment and miRNA regulation at AJs, since this can also occur through three LIM-domain containing actin-binding proteins, such as LMO7, LIMCH1, and PDLIM (*8*). We actually showed that all these components act on AGO2 through stabilization of the structure and tension of the apical actomyosin ring at AJs (*8*). Therefore, it is likely that disruption of either one of these components or of apical actomyosin can influence ECM remodeling. Since such disruption of AJs and of the epithelial barrier is common in diseases such as inflammatory bowel disease (IBD) and colon cancer (*1, 2, 91*), we predict that this may be a broader mechanism of ECM remodeling that can fuel these diseases. Although in this work we specifically examined PLEKHA7, it is likely that our studies describe a broader mechanism, which remains to be tested in future studies.

Due to their actomyosin connections, AJs are known mechanosensitive complexes that respond to mechanical cues, including those that stem from the ECM (*78, 92*). Along these lines, we have previously shown that collagen I and fibronectin negatively affect the localization and complex formation of PLEKHA7 and of the RNAi machinery at adherens junctions and throughout the cell (*93*). Since collagen I and fibronectin introduce an ECM that exerts stronger forces to cells, which can accordingly alter cytoskeletal dynamics (*21*), it is likely that the effects of these ECM components to the junctional RNAi machinery and miRNAs are mediated by the actin cytoskeleton, especially since we have shown that the latter is critical for AGO2 localization and miRNA function at AJs. However, this remains to be determined in upcoming studies. Still, taken altogether, disruption of PLEKHA7 and of the junctional miRNAs that results in ECM remodeling and collagen-fibronectin re-organization could in turn further disrupt the AJ-associated RNAi machinery, introducing a potential runaway feedback loop that further destabilizes miRNAs, accelerates ECM remodeling, and eventually alters cell behavior and tissue homeostasis. Indeed, PLEKHA7 KO cells plated in the stiff matrix (3 kPa) had lost both spherical shape and were larger, not only compared to the Caco2 WT cells plated on the same stiff matrix, but even compared to the same PLEKHA7 KO cells plated on the softer matrix (1.1 kPa), supporting this idea of an AJ-ECM feedback loop. These results outline a potential novel mode of interaction between AJs and ECM, through the RNAi machinery, which could influence epithelial cell behavior. Altogether, our recently published (*8, 93*) and present findings outline a mechanosensitive RNAi machinery that interacts with the ECM to fine-tune the mechanical responses and behavior of the cell. Fully delineating this crosstalk between the AJs and the ECM through RNAi and miRNAs and understanding all the components involved, can provide new insights into tissue homeostasis and disease pathogenesis, and will be the goal of our upcoming studies.

### Materials and Methods Cell culture

In all comparisons, cells were used at strictly the same confluences as specified for each experiment. All cell lines were authenticated by the University of Arizona Genetics Core (via Science Exchange) and checked for misidentified, cross-contaminated, or genetically drifted cells. Cell lines tested negative for mycoplasma contamination (LookOut Mycoplasma PCR Detection Kit-Sigma Aldrich). Human colon epithelial Caco2 cells (ATCC, Manassas-HTB-37) were grown in MEM cell culture medium (Corning-MT10010CV), supplemented with 10% FBS (Gibco-Life Technologies-A3160502), 1 mM sodium pyruvate (Gibco-Invitrogen-11360070-100 mM) and 1X non-essential amino-acid supplement (Gibco-Invitrogen-11140050-100x), at 37^0^C with 5% CO2. Human foreskin fibroblasts (HFF-1) were cultured in DMEM (Gibco-Fisher Scientific - SH3002301) supplemented with 10% FBS, 2 mM additional L-glutamine (Gibco -Thermo Fisher Scientific - 25030081), 1 mM sodium pyruvate, and 1X non-essential amino-acid supplement. Cells were maintained at 37°C with 5% CO2. For the ECM production, HFF-1 cells were seeded into 6 well plates for 2 days, when cell media were changed to media containing ascorbic acid for 4 more days. At day 6, HFF-1 cells were decellularized and Caco2 cells were seeded on top of the HFF-1 derived-ECM.

For the 3D RASTRUM culture experiments, 620,000 cells Caco2 WT and PLEKHA7-KO cells were plated using the RASTRUM bioprinter (Inventia Life Science) in 96 - well plates. Printing instructions for the machine were created using the online library of RASTRUM Cloud (Inventia Life Science). RASTRUM matrices used were PX02.28P - 1.1 kPa and PX03.28 - 3 kPa; activators used were F3, F177 and F178; Bioinks used were F32, F239 and F268. Cell culture media was changed every 48 hours and for 10 days.

For the 3D VitroGel® (TheWell Biosciences) culture experiments, cells were grown to 80% confluency in 100 mm cell culture dishes and transferred to 3D culture according to vendor instructions (The WellBiosceince SKU:VHM04 and SKU:VHM01) in 8-well chamber slides (ThermoFisher Scientific-Nunc™ Lab-Tek™ II-154534PK). In detail, 1x10^6^ cells per 1mL of cell culture medium with 30% FBS and 3X critical growth factors were prepared. Then the Vitrogel and the cell suspension were mixed at 2:1 ratio and 200 µL of the resulting hydrogel mixture was added to each well of the 8-well chamber slide. Gels were incubated at room temperature for 15 minutes. Then, 200 µL of additional fresh medium was each well added to cover the gels and incubated at 370C with 5% CO2 for 14 days. Covering medium was changed every 48 hours.

### Virus production and shRNA infection

Lentiviral shRNAs were derived from the pLKO.1-based TRC1 (Sigma–RNAi Consortium) shRNA library pLKO.1-puro; non-target shRNA control, SHC016; and PLEKHA7 shRNA (shPLEKHA7), TRCN0000146289. Lentiviruses were produced in HEK 293FT cells and used to infect cells according to standard protocols. In detail, lentiviral particles were produced in HEK 293FT cells co-transfected via Lipofectamine 2000 (Invitrogen-ThermoFisher Scientific- 11668019) with the 4 µg of shRNA vector and 3.5 µg of Virapower packaging plasmids (pMDLg/pRRE, Addgene-12251; pRSV-Rev, Addgene-12253; pMD2.G, Addgene-12259) according to manufacturer instructions. shRNA lentivirus was collected and used to infect Caco2 cells. More specifically, Caco2 cells were grown to 40% confluency in 60mm cell culture dishes, washed with 1X PBS once, and then 1.5 mL of fresh medium including 8 µg/mL of polybrene (Sigma-Aldrich: #H9268) was added to each plate and incubated at room temperature for five minutes. Afterwards, 500 µL of lentivirus was added to the cells and grown for 24 h at 37^0^C with 5% CO2. After 24 hours, fresh medium was added to the cells and were grown for another 48 hours before selecting with 5 µg/mL puromycin (Gibco-ThermoFisher Scientific-A1113803) for 72 hours. Stable shRNA-expressing cell lines were maintained in 2.5 μg/ml puromycin following initial selection.

### CRISPR/Cas9 deletion of *PLEKHA7* in Caco2 cells

The PLEKHA7 knockout Caco2 cell line was generated using CRISPR/Cas9 technology and previously published (*8*). The sgRNA sequence (GGATGTTTCACTGACCATGCTGG) targeting exon 5 of PLEKHA7 was designed using the online Synthego tool (www.synthego.com) with the help of the Cell Models Core (COBRE in Digestive and Liver Diseases, MUSC) and synthesized by Integrated DNA Technologies, IDT. This PLEKHA7-targeting sgRNA was cloned into the PX459 pSPCas9(BB)-2A-Puro vector. Caco2 cells were transfected with 30 µg the sgRNA-containing vector using Lipofectamine 3000 (Invitrogen-ThermoFisher Scientific - L3000001) according to the manufacturer’s protocol. After 24 hours, cells were selected with 5 μg/ml puromycin for 48 hrs. Successfully transfected cells were seeded to obtain single colonies, which were picked and expanded for screening. PCR using primers designed to amplify the target region (Forward: CCTTTCTCAGGGCCTCATGTCA; Reverse: GCCTCCAAACAATCAGGGTTGG) was performed with DNA extracted from colonies using QuickExtract (Lucigen-ThermoFisher Scientific- QE0905T). PCR amplified products were screened by restriction digest analysis using the CviAII restriction enzyme (New England Biolabs - R0640S). Positive candidates were sequenced to identify and confirm indels. See also (*8*).

### miRNA mimic transfections

miRNA mimic transfections were conducted using Lipofectamine™ RNAiMAX Transfection Reagent (ThermoFisher Scientific – 13778075) according to the manufacturer’s guidelines. In detail, Caco2 wild type and *PLEKHA7* knockout cells were grown at 60% confluency in 6-well plates. Then, 150 µL of Opti-MEM™ - Reduced Serum Medium (Gibco ™ - 31985062) was mixed with 9 µL of RNAiMAX reagent, whereas in a separate tube, 150 µL of Opti-MEM™ - Reduced Serum Medium was mixed with desired miRNA mimic to obtain a final concentration of 25 pmol. The solutions containing the RNAiMAX reagent, and the miRNA mimic were mixed and incubated at room temperature for 15 minutes. The above amounts are for one well of a 6-well plate. In the meantime, cells were washed with 1X PBS once and replaced with 1.7 mL of fresh medium per well. After the incubation was completed, 300 µL of miRNA-lipid complex was added to the cells and grown for 24 h at 37 ^0^C with 5% CO2. After 24 hours, fresh medium was added to the cells and were grown for another 48 hours. Cells were then collected to analyze protein and/or RNA. MISSION miRNA mimics (Sigma Aldrich) used: hsa-miR-203a, HMI0357-5NMOL; HMI0456- 5NMOL; hsa-miR-30c-5p, HMI0460-5NMOL; hsa-miR-24-3p, HMI0410-5NMOL; hsa-let-7g, HMI0020-5NMOL; negative control miRNA mimic, HMC0002-5NMOL.

### Luciferase constructs and assay

Constructs for performing luciferase-based assays were designed using the Snapgene software tool (www.snapgene.com), the miRbBase database (release 22.1)(*94*), the NCBI BLAST tool, and the NCBI reference sequences of 3’UTR of LOX (NM_002317.7) and MEP1A (XM_011514628.2). 3’ UTRs of LOX and MMP1 with and without mutations were cloned to the pmirGLO Dual-Luciferase miRNA target expression vector (Promega-E1330) by GeneScript vector construction services (www.genescript.com). Luciferase assays were conducted using previously published methods (*95*) and vendor instructions (Lipofectamine™ 3000 transfection reagent – Invitrogen- L3000001). In detail, cells were grown to 90% confluency in 96-well, white-bottom plates (Fisher Scientific-Corning-353296). Then, 0.15 µL of Lipofectamine 3000 reagent was mixed with 5 µL of OPTI-MEM medium in a 1.5 mL tube. In a separate tube, 5 µL of OPTI-MEM medium was mixed with 0.2 µL of P3000 reagent, 0.1 µg of luciferase construct, and 30 pmol of miRNA mimic (all calculations are for one well of 96 well plate). Then, diluted luciferase construct and miRNA mimic was added to the diluted Lipofectamine 3000 and incubated at room temperature for 20 minutes. 65 µL of fresh cell culture medium was added to each well with 10 µL of plasmid-miRNA-lipid complex to reach a total volume of 75 µL and incubated at 37^0^C with 5% CO_2_ for 48 hours. After 48 hours, an additional 75 µL of fresh medium was added to each well. Luciferase activity was measured using Dual-Glo® Luciferase Assay System (Promega-E2920) per vendor instructions. In detail, 75 µL of Dual-Glo reagent was added to each well and incubated at room temperature for 30 minutes to measure firefly luminescence. Next, 75 µL of Dual-Glo-Stop & Glo reagent was added to each well and incubated at room temperature for 30 minutes, followed by renilla luminescence measurements. The ratio between the firefly luminescence and the renilla luminescence was calculated and normalized for the control construct. Luminescence measurements were taken using a Synergy H1 Hybrid Multi-Mode Readers. Sequences of miRNA mimics: hsa-miR-24-3p, 5’-UGGCUCAGUUCAGCAGGAACAG-3’; hsa-miR-30c-5p, 5’- UGUAAACAUCCUACACUCUCAGC-3’. Primers used: LOX 3’UTR: forward: 5’- TTGGAGCTCCCGCTCTCCTCCTTCCTTCAC-3’; reverse: 5’-TGGCTCGAGTTGGACAAATTTCCAGTGTGACC-3’. The seed sequence of hsa-miR-24-3p for LOX 5’-ACTGAGCCA-3’ was mutated to 5’-ACTAATCAA-3’. The seed sequences for of hsa-miR-30c-5p for LOX 5’-ATGTTTACA-3’ mutated to 5’-TTGATAA-3’, 5’-GTAGGAT-3’ mutated to 5’-TTAAGTT-3’, 5’-GCTGAGA-3’ mutated to 5’-GATAATA-3’, 5’-GTTTACA-3’mutated to 5’-TTGTGCA-3’, 5’-CTGAGAGT-3’ mutated to 5’-CCGATATA-3’.

### RNA isolation, RNA sequencing and analysis, qRT-PCR

Caco2 cells were lysed using Trizol (Invitrogen) and subjected to the Trizol Plus total transcriptome isolation protocol of the PureLink RNA mini kit (Ambion, Life Technologies) specified to isolate both mRNAs and miRNAs. RNA concentrations were determined using a BioDrop uLite spectrophotometer (Harvard BioScience). For sequencing, 100–200 ng of total RNA was used to prepare RNA-Seq libraries using the TruSeq RNA Sample Prep Kit (Illumina), following the protocol described by the manufacturer. High-throughput sequencing (HTS) was performed using an Illumina HiSeq2500 with each sample sequenced to a minimum depth of ∼ 50 million reads. Illumina Casava1.8 software used for basecalling. Sequenced reads (fastq files) were trimmed for adaptor sequence and masked for low-complexity or low-quality sequence. Secondary analysis was carried out on an OnRamp Bioinformatics Genomics Research Platform (OnRamp Bioinformatics, San Diego, CA). OnRamp’s advanced Genomics Analysis Engine utilized an automated RNAseq workflow to process the data, including data validation and quality control and read alignment to the human genome (hg19) using blastx. The resulting SAM files were sorted and inputted into the Python package HTSeq to generate count data for gene-level differential expression analyses. Transcript count data from DESeq2 analysis of the samples were sorted according to their adjusted p-value or q-value, which is the smallest false discovery rate (FDR) at which a transcript is called significant. Sequencing and analysis was performed by Bob Wilson, Willian Da Silveira, Hazard Starr, and Gary Hardiman, at MUSC Genomics and Bioinformatics Cores. Data are deposited in the GEO public database, under the accession number GSE156860. For qRT-PCR, RNA was converted to cDNA using the High Capacity cDNA Reverse Transcriptase Kit (Applied Biosystems). qPCR reactions were performed using the Taqman FAST Universal PCR master mix (Applied Biosystems), in a BioRad CFX96 Touch/Connect real time quantitative PCR machine. TaqMan assays used - Applied Biosystems, cat# 4427975: hsa-miR-24, 000402; hsa-miR-30c, 000419; U6, 001973; Applied Biosystems cat# 4331182: human 18S, Hs99999901_s1; mouse 18S, Mm04277571_s1; Lox: Mm00495386_m1; Applied Biosystems cat# 4448892: Plekha7, Mm00555062_m1; Jun, Mm007296811_m1.

### Pathway and bioinformatic analysis

The mRNA data from the analysis were ranked according to the highest statistical significance of - log_10_ (p-value) and highest differential expression according to fold change. mRNAs that were increased upon PLEKHA7 depletion by more than or equal to 1.5-fold were subjected to pathway analysis using online tool Reactome database, which includes 14,628 human reactions, 2,629 pathways and 11,396 different human genes (*32*). The entities ratio is the fraction of the 498 mRNAs (“entities”) matched within a specific pathway in the dataset against the total number of genes in that specific pathway in the database (“entities total”). The “reaction ratio” is the fraction of identified sub-pathways against the entire biological interactions in the database (Fig. 1B, table S2, S3). The list of mRNAs with ≥ 1.5 fold change was then compared to the Matrisome database (*33, 34*). PLEKHA7-enriched miRNAs from our previously published research (*9–11*) were tested against the selected ECM associated mRNAs using online tools DIANA TOOLS - version 8.0 (*42*), mirTarBase - release 8.0 (*43*), and TargetScan – releases 7.2 and 8.0 (*44, 45*) to predict targeting possibilities.

### Immunoblotting

Whole cell or mouse tissue extracts were obtained using RIPA buffer (50 mM Tris pH 7.4, BioRad 1610719; 150 mM NaCl, Sigma Aldrich S9888-10K; 1% NP-40, Fisher Scientific 507517565; 0.5% deoxycholic acid, Sigma Aldrich D6750-100G; and 0.1% SDS, Thermo Fisher Scientific, BP166-500), supplemented with protease (Thermo Fisher Scientific, 50550432) and phosphatase (Pierce Biotechnology, P178420) inhibitors. Lysates were homogenized by passing through a 29-G needle and cleared by full-speed centrifugation for 5 min. Protein quantification was performed using a Pierce BCA Protein Assay (Pierce Biotechnology, I23227). Protein extracts were mixed with Laemmli sample buffer at 2x final, separated by SDS-PAGE using 4-20% TGX gels (Bio-Rad, 4568094), and transferred to 0.2 m nitrocellulose membranes (Bio-Rad, 1704158) with the Bio-Rad® Trans-Blot Turbo Transfer System. Membranes were blocked and blotted in 3% milk according to standard protocols. Antibodies used: PLEKHA7, Sigma Aldrich HPA038610 (1:1000 dilution); MEP1A, Invitrogen PA5-28371 (1:2000 dilution); MMP1 E9S9N, Cell Signaling 54376S (1:500 dilution); LOX, Abcam ab31238 (1:500 dilution); JUN, Cell Signaling 9165 (1:1000 dilution); GAPDH, Cell Signaling Technology 14C10-2118 (1:2000 dilution); and Actin, Cell Signaling Technology 4967L (1:2000 dilution). Signals were detected by luminescence using Pierce ECL (Bio-Rad, 32209) using a Bio-Rad® ChemiDoc Imaging System.

### LOX and MMP inhibition and activity measurements

Cells were grown to 80% confluency in 100 mm cell culture dishes and treated with either Lysyl Oxidase (LOX) inhibitor BAPN (3-Aminopropionitrile fumarate salt; Millipore Sigma -A3134-5G, at 150 µM final concentration, or the MMP inhibitor Doxycycline Hyclate CAS 24390-14-5 (Santa Cruz-sc-204734B) at 0.2 µg/mL final concentration, for 24 hours before collecting protein or performing assays. The working concentrations of the inhibitors used were based on prior research (*96, 97*). For the LOX activity assay, cells were washed in cold 1X PBS once, then 100 µL of cold 1X PBS was added to the cells and scrapped. Cells were homogenized quickly by pipetting up and down and the homogenate was spun down in at 13,000g, at 4°C, for 5 mins. The supernatant was used to measure the fluorescence activity according to vendor’s instructions (LOX activity; Abcam, ab112139; Abcam, ab284573) at Ex/Em = 530-570/590-600 nm, at 37°C. For the MMP/collagenase activity assay, cells were washed with cold 1X PBS once, and homogenized in 100 µL of the provided lysis buffer (Abcam, ab234624) on ice, for 5 mins. Homogenates wer spun down at 16,000g at 4°C, for 10 mins. The supernatants were used to measure fluorescence activity according to vendor’s instructions (Abcam, ab234624) at Ex/Em = 490/520 nm, at 37°C.

### Cell invasion assay

This assay was conducted using the xCELLigence RTCA CIM plate 16 (xCELLigence, 5665817001) on the xCELLigence RTCA DP instrument, according to vendor’s instructions. In detail, the upper chamber of the RTCA CIM plate 16 was coated with 50 µL of 10 µg/mL collagen-I to each well and incubated at 37^0^C for 2 hours. Afterwards, excess collagen-I was removed, and wells were washed once with 1X PBS. Then, 160 µl of Caco2 cell culture medium containing 10% FBS was added to each well of the bottom chamber, while 50 µl of the Caco2 cell culture medium containing 2% FBS was added to the wells of the top chamber. After assembling the top and bottom chamber, the CIM-16 plate was left at 37^0^C incubator for one hour to obtain background measurements. Then, 30,000 cells in 100 µL of medium per well and of each condition were added to the top chamber and the CIM-16 plate was left at room temperature for 30 minutes. Thereafter, the CIM-16 plate was loaded to the RTCA DP instrument and the measurements were taken at every 15 minutes for 48 hours.

### Cell migration assay

Cells were grown to 80% confluency on Matrigel (Corning-ThermoFisher Scientific, CB40230C) coated coverslips. Next, 2-well inserts (Ibidi, 80209) were placed on Matrigel coated coverslips. 17500 cells in 70 µL of cell culture medium was placed in each well of the inserts and were grown for 24 hours to full confluency at 37 ^0^C with 5% CO2. After 24 hours, 2-well inserts were removed from the coverslips and fresh medium was added. The time point of the 2-well inserts removal was time point zero for quantification of cell migration. Images of cell migration were taken every 10 hours for 30 hours. Experiment was stopped at 30 hours, to avoid any cell proliferation effects that would influence the results, since Caco2 cells have a doubling time of 32 hours (*60*).

### Total collagen assay

Total Collagen Assay (Perchlorate-Free; Abcam, ab222942) was performed to detect insoluble collagen per the manufacturer’s instructions. Briefly, HFF-1 cells were seeded into 6 well plate and 2 days later media of cells were changed with media containing ascorbic acid for 4 more days. At day 6, HFF-1 cells were decellularized and Caco2 cells were seeded on top of the HFF-1 - derived ECM. 48 hours later, cells and modified ECM were solubilized in acid/pepsin mixture (0.5 M acetic acid and 0.1 mg/ml of pepsin) overnight at 4°C. The insoluble crosslinked collagen was collected after centrifugation at 10000g for 20 minutes. Finally, the insoluble pellet was dissolved in 10 N NaOH at 120 °C for 1 hour followed by neutralization with 10 N HCl. Total collagen was detected via the kit’s reagent.

### Decellularized assays and immunofluorescence staining

HFF-1 cells were seeded on µ-Slide 8-well (Ibidi, Germany) glass for immunofluorescence staining and 2 days later, media of cells were changed with media containing ascorbic acid for 4 more days. At day 6, HFF-1 cells were decellularized using an extraction buffer containing 25 mM ammonium hydroxide and 0.5% Triton X-100; 20 ug/ml DNAse I treatment was done for 30 min to get rid of the remaining cell debris (*98*). Then, Caco2 wild type or PELKHA7 KO cells were seeded on top of the HFF-1 - derived ECM. 48 hours later, cells were fixed with 4% paraformaldehyde for 15 min, permeabilized and blocked in 3% BSA-PBS. Cells were incubated in Type I collagen (Abcam, ab34710) and fibronectin (Santa Cruz, sc-29011) primary antibodies overnight at 4°C and secondary Alexa Fluor 594 or 488-labeled antibodies for 2 hours at room temperature. Cells were also counterstained with DAPI for 5 min at room temperature. Images were acquired using a ZEN Black LSM 880 Airyscan (Carl Zeiss, DE). Image ECM quantifications were performed using the TOWMBLI macro on Fiji (*54*).

### Animal approval, generation of mutant mice, genotyping, tissue collection and tissue analysis

All experiments on mice were approved by the Institutional Animal Care and Use Committee of the Medical University of South Carolina and the MUSC Transgenic Mouse Core Facility under the protocol numbers IACUC-2018-00280 and IACUC-2021-01198. The *Plekha7* mutants were generated directly on an inbred C57BL/6J strain background by CRISPR/Cas9-mediated genome editing that directly targeted the zygotes of C57/BL6J mice. We deleted the genomic sequence between exons 2-10 to eliminate all possible Plekha7 isoforms using guide RNAs 5’- ggtgcactctcactcaactcagg-3’ (MS1042.sp2) and 5’-gacgtgacaccgccaggcaatgg-3’ (MS1042.sp6) (Fig. S4). The mouse model was generated through MUSC’s Transgenics Core and Dr. Alexander Awgulewitsch. Whole genome sequencing in DNA from knockout animals was performed to confirm that there were no other mutations or deletions across the genome, other than the *Plekha7* deletion. Male and female mice of ≥6 weeks of age were used for breeding. Animals were kept in a 12-hour light-dark cycle with food and water ad libitum. Genetic drift was avoided through inbreeding of heterozygotes, after F1. For genotyping, DNA and protein were isolated from tail clips or ear punches at weaning on 21^st^ day after putting animals under anesthesia by 2-4% Isoflurane inhalation. Animals were euthanized by CO_2_ gas, followed by cervical dislocation prior to tissue collection. Approximately 50 mm long colon tissues were obtained near the caecum (proximal colon) and near anus (distal colon). Genotyping was conducted using Phire Tissue Direct PCR Master Mix (ThermoFisher Scientific- F170S) according to the vendor’s instructions. Primers used were, forward: 5’-AAGAGCTAGCCCTGAAACCTG-3’, reverse: 5’- TCTGCCTAGAGGATGGTGCT-3’, 5’-ACTGGTGAGGGAAAGGGTTT-3’.

For RNA isolation, tissues were lysed using Trizol (Invitrogen) and subjected to the Trizol Plus total transcriptome isolation protocol of the PureLink RNA mini kit (Ambion, Life Technologies) specified to isolate both mRNAs and miRNAs, as described above. For tissue stainings, mouse colon tissues were fixed in 10% formalin, embedded in paraffin, and sectioned in slides. To deparaffinize, slides were immersed in xylene (Fisher Scientific) twice for 5 min each time and were rehydrated through a series of EtOH (100-100-95-80-70-50%) and placed in water. (a) For H&E staining, slides were stained with hematoxylin (Polysciences, cat # 24243-500) for 2 min and washed in water-acid alcohol-water. Slides were then immersed in blue ammonia water for 1 min, washed with water, placed in 95% EtOH for 1 min and stained with in eosin Y solution (Fisher Scientific, cat # 45380-M1159350100) for 1 min. Slides were then rapidly dehydrated through a series of EtOH (95-95-100-100%), placed in xylenes twice for 5 min each time and were mounted with Cytoseal (Fisher Scientific, cat # 831016). Images were obtained using Keyence fluorescence microscope under 20X - Plan Apochromat 20X BZ-PA20-VHX-E20 objective lens (Keyence) using the tissue perimeter detection method. (b) For immunofluorescence, deparaffinized slides were subjected to antigen retrieval for 32 mins at 95 °C with EDTA and incubated with the respected antibodies for 32 mins at 37 °C using a Ventana Discovery Ultra system (Roche). Antibodies used were PLEKHA7, Genetex (cat # GTX131146) at 1: 200 dilution and E-cadherin Cell Signaling Technologies (cat #3195) at 1:300 dilution. Slides were scanned using an Akoya Vectra Polaris scanner and images were analyzed using QuPath 0.2.0.

### Image acquisition, analysis, and quantifications

For second-harmonic generation, images were taken under Plan Fluorite 20X LD PH BZ-PF20LP (Keyence) objective lens with Jigtech Polarizer-N11599-00R-002 (Keyence) to polarize the light using the tissue perimeter detection method. To analyze second harmonic generation images, the average fluorescence from the entire tissue of one animal and the average fluorescence in the basement membrane area of three different fields of the same tissue from each animal were measured using Fiji/ImageJ software. In addition, basement membrane area with fluorescence of three different fields of the same tissue from one animal was measured. For the cell migration assay, images were taken using a Leica-DM1000 microscope at 20X magnification. Images were uploaded to Fiji/ImageJ software, and the area of the cell growth was measured for each condition. For each time point a minimum of 3 and a maximum of 6 images were taken. For statistical comparisons, the area of growth between Caco2 cells and PLEKHA7 depleted Caco2 cells at each time point was measured. Western blot quantifications were performed using Fiji/ImageJ software (*99*).

### Statistical analysis

Unless otherwise stated, all error bars are shown as means ± SD, and a student’s t-test with p < 0.05 was used to detect statistically significant differences between two groups. P-values are listed in the figure legend of each image. Graphs were generated using SPSS v24, SPSS v25, Microsoft Excel, GraphPad Prism9, or R studio. Boxplots depict data quartiles and the error bars on these graphs represent the 95% confidence interval. To analyze results from the ECM combination experiments and all the other experiments with three or more groups one-way ANOVA was utilized, where we used Shapiro-Wilk test for the normality test and Bonferroni correction with p<0.05 for pair-wise comparisons. In all mage analysis, a scale was set to keep the pixel/micron ratio constant. Number of replicates are listed in the figure legend of each image.

### Supplementary Materials

Figs. S1 to S4 Tables S1 to S6

## Supporting information

Daulagala et al - SupplTables S1-6

## Acknowledgments

We would like to thank Bob Wilson, Willian Da Silveira, Hazard Starr, and Gary Hardiman at MUSC Genomics and Bioinformatics Cores for RNAseq and analysis; MUSC’s Transgenics Core and Dr. Alexander Awgulewitsch for mouse knockout generation; MUSC’s Division of Lab Animal Resources (DLAR) and Dr. Kris Helke, for help with animal handling and tissue extraction; Drs. Christiana Kappler and Stephen Duncan, Cell Models Core, MUSC Center of Biomedical Research Excellence (COBRE) in Digestive & Liver Disease (CDLD; NIH P20 GM130457) and MUSC Digestive Disease Research Center (DDRC; NIH P30 DK123704) for support with CRISPR/Cas9 gene knockouts; the Ralph Johnson VA Medical Center imaging core for help with second harmonic generation imaging; the Translational Science Lab (Melodie Parrish, Elizabeth O’Quinn) Hollings Cancer Center, MUSC (NIH P30 CA138313) for tissue immunofluorescence; the shRNA Technology Shared Resource (David Turner), Hollings Cancer Center, MUSC (NIH P30 CA138313) for shRNA constructs; the Molecular Analytics Core, Department of Regenerative Medicine and Cell Biology and Dr. Jamie Barth for qRT-PCR instrumentation support; and Dr. Mindy Engevik’s laboratory, for allowing us to use their Synergy H1 Hybrid Multi-Mode Reader.

## Funding

This work was supported by NIH grants R01 DK124553, R01 DK136658, 3P20 P20 GM130457 (COBRE in Digestive & Liver Disease, MUSC; GM130457-04S1 supplement), P30 DK123704 (Digestive Disease Research Center, MUSC), Concern Foundation (Conquer Cancer Now Award), and P30 CA138313 (Hollings Cancer Center Pre-Clinical & Clinical Concepts Pilot Award) to AK. AD was supported by the Abney Graduate Fellowship Award, Hollings Cancer Center (P30 CA138313), MUSC. MCB was supported by NIH training grants TL1 TR001451 and UL1 TR001450. MC and OS were supported in part by R01CA267101. MC is supported by the Hollings Cancer Center Postdoctoral Fellowship.; AB - support. ADB was supported by Dept. of Veterans Affairs.

## Author contributions

Conceptualization: AD, AK; Methodology: AD, MC, JNM, DWJ, MCB, AB, OS, AK; Investigation: AD, MC, JNM, DWJ, MCB, AB, OS, AK; Visualization: AD, MC, JNM, AK; Supervision: AK, OS; Writing—original draft: AD, AK; Writing—review & editing: AD, MC, JNM, DWJ, MCB, AB, OS, AK.

## Competing interests

OS is the founder and the president of LoxiGen, Inc, developing LOX inhibitors. The other authors declare that they have no competing interests.

## Data and materials availability

All data are available in the main text, figures, or the supplementary materials. Sequencing data have been deposited in public databases and accession numbers are provided in the related sections within the text.

---

**Fig. S1.**
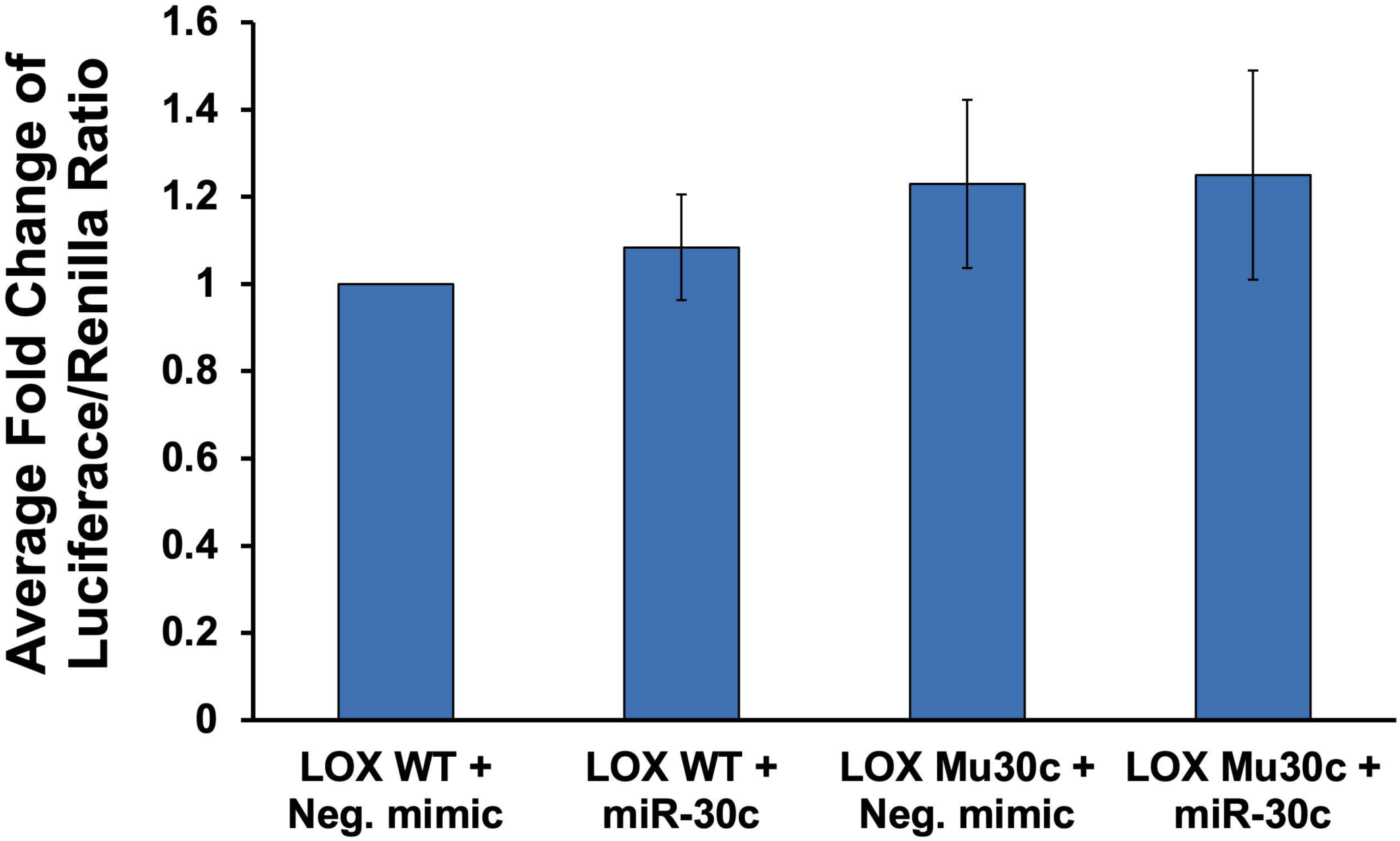
miR-30c does not target LOX mRNA directly. Luciferase reporter assay using LOX 3’UTR luciferase constructs, wild type (LOX WT) and mutated for the miR-30c target sequence (LOX Mu30c), transfected in PLEKHA7 KO Caco2 cells and co-transfected with negative control miRNA mimic, or miR-30c mimic. Results shown are the average luciferase/renilla luminescence ratio from n=3 biological replicates.

**Fig. S2.**
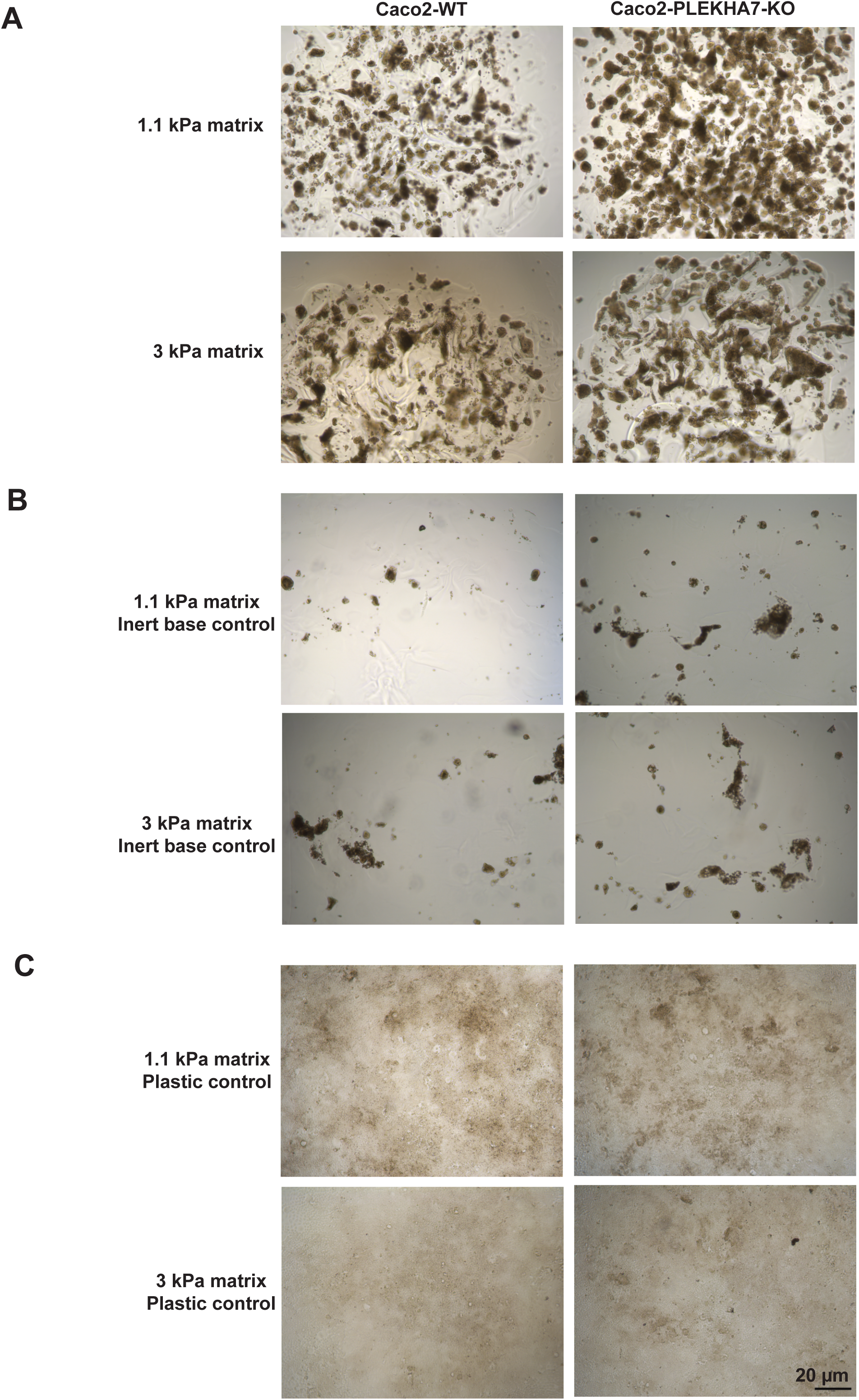
PLEKHA7-KO Caco2 cells exhibit abnormal growth in stiff matrices. **(A)** Wider representative fields of Caco2-WT and PLEKHA7-KO cells that were grown in 3D cultures in 1.1 kPa and in stiffer, 3 kPa matrices, using a RASTRUM apparatus for 10 days, as in Fig. 7. (**B**) In all conditions, there were significantly less colonies in the inert base control, which only had the first layer of coating and did not have the matrix with the synthetic ECM peptides. (**C**) In all plastic- only controls, which did not have either layer of coating or the matrix with the synthetic ECM peptides, cells did not grow in spheroids.

**Fig. S3.**
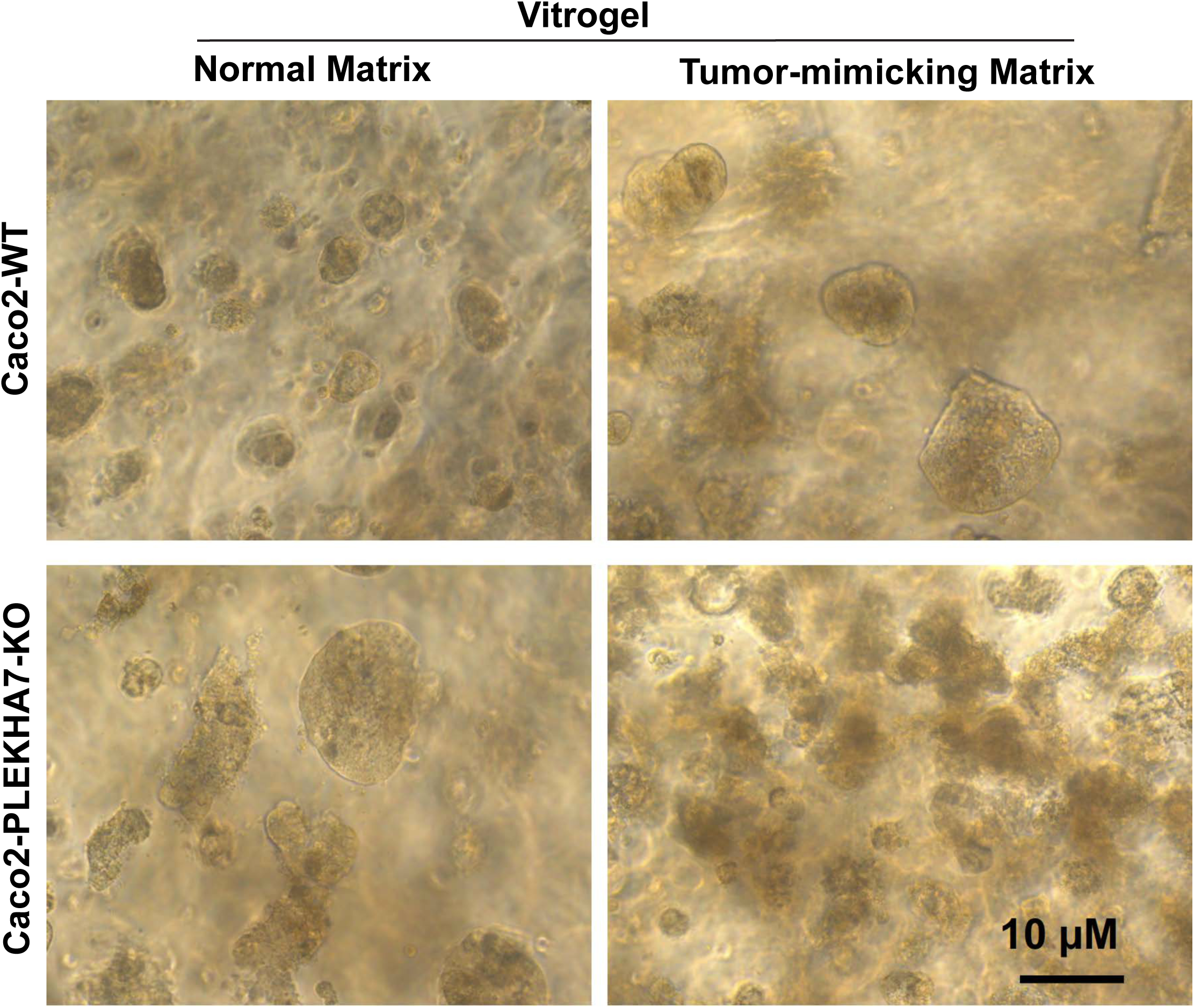
PLEKHA7-KO cells grow irregularly in stiff, tumor-mimicking matrices. Caco2-WT and PLEKHA7 KO cells were grown on hydrogels (Vitrogel) that mimic normal and stiffer, tumor- mimicking matrices for 14 days. Results resemble those obtained using the RASTRUM 3D cultures (Fig. 7 and S2) and show progressive increase in size and loss of circularity of PLEKHA7- KO cells and in the stiffer matrices.

**Fig. S4.**
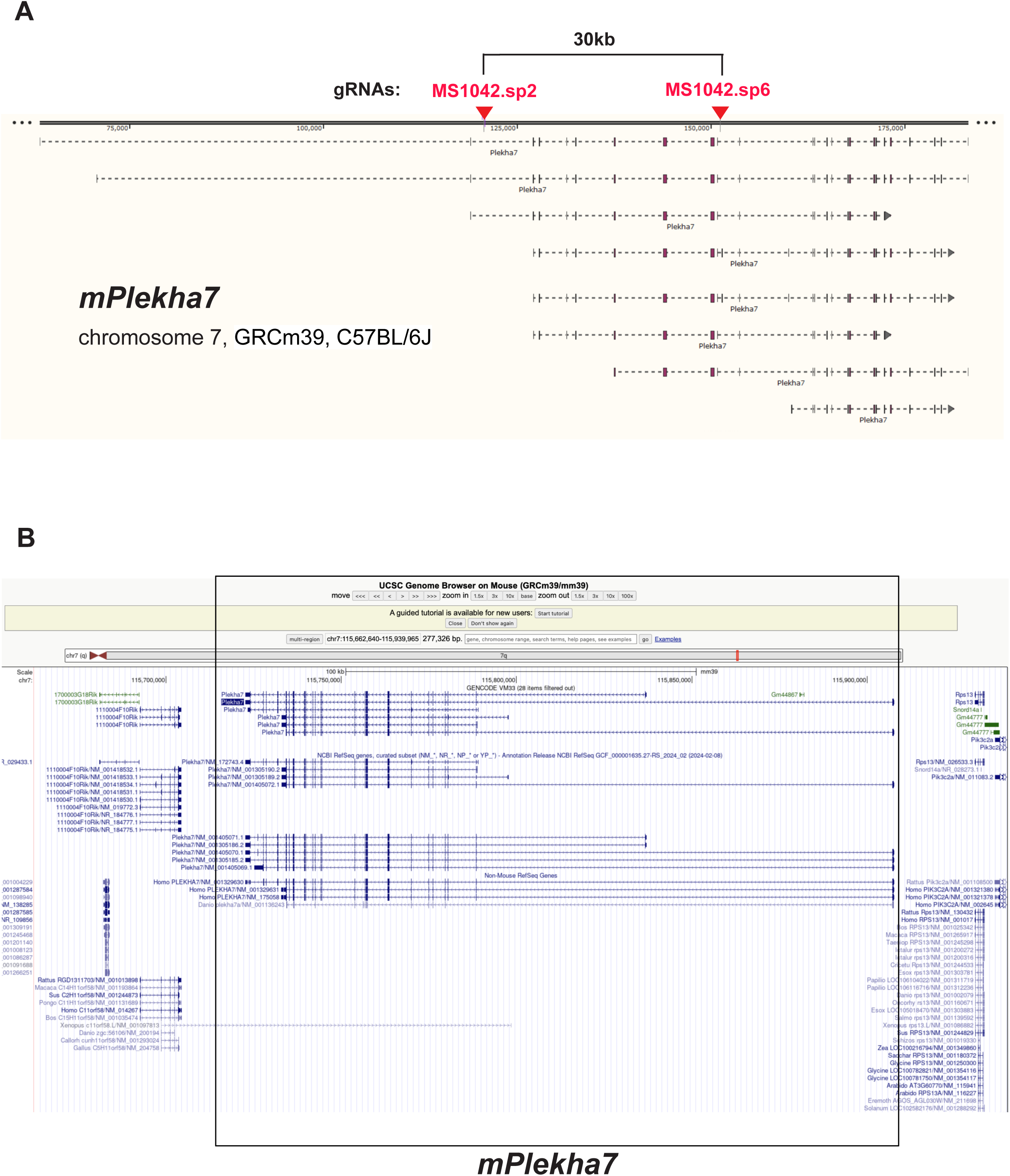
Generation of *Plekha7* knockout mice. **(A)** Outline of the mouse *Plekha7* genomic region, indicating the gRNAs used and their targeting positions to cleave a 30 kb region between exons 2- 10 using CRISPR/Cas9, as well as the different *Plekha7* mRNA isoforms predicted to be expressed. **(B)** Screenshot of the UCSC mouse genome browser (mm39) outlining the mouse *Plekha7* genomic region.

